# A Timer for analyzing temporally dynamic changes in transcription during differentiation in vivo

**DOI:** 10.1101/217687

**Authors:** David Bending, Paz Prieto Martín, Alina Paduraru, Catherine Ducker, Erik Marzaganov, Marie Laviron, Satsuki Kitano, Hitoshi Miyachi, Tessa Crompton, Masahiro Ono

**Affiliations:** Department of Life Sciences, Faculty of Natural Sciences, Imperial College London, Sir Alexander Fleming Building, Imperial College Road, London, SW7 2AZ, United Kingdom; Institute for Viral Research, Kyoto University, Sakyo-ku, Kyoto 606 8507; UCL Great Ormond Street Institute of Child Health, 30 Guilford Street, London, WC1N 1EH, United Kingdom

**Author notes:** Present address: Institute of Immunology and Immunotherapy, College of Medical and Dental Sciences, University of Birmingham, B15 2TT, United Kingdom (David Bending). Correspondence author: Dr Masahiro Ono, Department of Life Sciences, Sir Alexander Fleming Building, Imperial College Road, South Kensington, London, SW7 2AZ, United Kingdom.

## Abstract

Understanding the mechanisms of cellular differentiation is challenging because differentiation is initiated by signaling pathways that drive temporally dynamic processes, which are difficult to analyse in vivo. We establish a new Tool, Timer-of-cell-kinetics-and-activity (Tocky [toki], time in Japanese). Tocky uses the Fluorescent Timer protein, which spontaneously shifts its emission spectrum from blue-to-red, in combination with computer algorithms to reveal the dynamics of differentiation in vivo. Using a transcriptional target of T cell receptor (TCR)-signaling, we establish Nr4a3-Tocky to follow downstream effects of TCR signaling. Nr4a3-Tocky reveals the temporal sequence of events during regulatory T cell (Treg) differentiation and shows that persistent TCR signals occur during Treg generation. Remarkably, antigen-specific T cells at the site of autoimmune inflammation also show persistent TCR signaling. In addition, by generating Foxp3-Tocky, we reveal the in vivo dynamics of demethylation of the *Foxp3* gene. Thus, Tocky is a Tool for cell biologists to address previously inaccessible questions by directly revealing dynamic processes in vivo.

**Summary:** The authors establish a new Tool, Timer-of-cell-kinetics-and-activity (Tocky) revealing the temporal dynamics of cellular activation and differentiation in vivo. The tool analyses the temporal sequence of molecular processes during cellular differentiation and identifies cells that receive persistent signals in vivo.

## Introduction

It is a central question in cell biology how cellular differentiation progressively occurs through the activities of temporally-coordinated molecular mechanisms (Kohwi and Doe, 2013; Kurd and Robey, 2016). It is, however, challenging to investigate in vivo mechanisms at the single cell level because individual cells are not synchronized and are heterogeneous, receiving key signalling at different times and frequencies in the body. No existing technologies can systematically analyse the temporal dynamics of differentiation and activities of individual cells in vivo. Intravital microscopy is useful for analysing cells in micro-environments (Koechlein et al., 2016) but is not suitable for systematically analysing cells which rapidly migrate through tissues, such as T cells. Single cell sequencing can provide ‘pseudotime’, but this is not the measurement of time, as the name implies, but is a measurement of the transcriptional similarities between samples at chosen analysis time points (Trapnell et al., 2014). Flow cytometry is suitable for determining the differentiation stage of individual cells, but current methods cannot be applied to investigate how individual cells sequentially differentiate into more mature stages, as data from individual cells do not currently encode time information (Hoppe et al., 2014). There is thus a great need for a new technology to experimentally analyse the passage of time following a key differentiation event, or *the time domain*, of individual cells in vivo. Such a new technology would benefit all areas of cellular biology, but would be particularly useful for the study of T-cells under physiological conditions in vivo, where both the time and frequency of signaling are critical to their differentiation.

T cells migrate through the body (Krummel et al., 2016), and their activation and differentiation statuses are almost exclusively determined by flow cytometric analysis (Fujii et al., 2016). In T cells, T cell receptor (TCR) signaling triggers their activation and differentiation (Cantrell, 2015), and is the central determinant of thymic T cell selection (Kurd and Robey, 2016) (including negative selection (Stepanek et al., 2014) and Treg selection (Picca et al., 2006)) and antigen recognition in the periphery (Cantrell, 2015). While the temporal dynamics of proximal TCR signaling, which are in the time scale of seconds, have been comprehensively and quantitatively analysed (Roncagalli et al., 2014; Stepanek et al., 2014), it is still unclear how transcriptional mechanisms for activation and differentiation respond to TCR signals over time in vivo. Such a transcriptional mechanism may be used for a new reporter system to analyse the dynamics of T cell activation and differentiation upon antigen recognition.

TCR signalling activates NFAT, AP-1 and NF-κB, which initiate the transcription of immediate early genes within a few hours (Oh and Ghosh, 2013), but their effects on T cell differentiation over the time scale of hours and days are obscure. In order to analyse TCR signal strength, currently, *Nur77 (Nr4a1)-EGFP* reporter mouse is commonly used (Moran et al., 2011), but the long half-life of the reporter gene, EGFP (~ 56 h (Sacchetti et al., 2001)) prevents its application for the analysis of the temporal dynamics of the events downstream of TCR signaling in vivo.

In this study, we have established a new Fluorescent Timer technology, **T**imer **o**f **C**ell **K**inetics and Activit**y** (Tocky), which uniquely reveals the time and frequency domains of cellular differentiation and function in vivo. Fluorescent Timer proteins have been used to analyse in vivo protein dynamics and receptor turnover (Dona et al., 2013; Khmelinskii et al., 2012), as well as identify progenitor cells, i.e. those cells expressing only immature fluorescence during embryogenesis and pancreatic beta cell development (Miyatsuka et al., 2011; Miyatsuka et al., 2014; Subach et al., 2009; Terskikh et al., 2000). However, those studies were qualitative and did not recognise the quantitative power of Fluorescent Timer. Here we develop a new Fluorescent Timer approach to quantitatively analyse the time and frequency domains of gene transcription within individual cells in vivo. By identifying a downstream gene of TCR signalling *(Nr4a3)* and developing a Fluorescent Timer reporter mice for the gene, we experimentally establish and validate the Tocky system for TCR signalling. Furthermore, we apply the Tocky approach to the *Foxp3* gene, which is the lineage-specific transcription factor of regulatory T cells (Treg), revealing in vivo dynamics of Treg differentiation. Thus, Tocky technology reveals time-dependent mechanisms of in vivo cellular differentiation and developmental states following key signalling pathway or lineage commitment, which cannot be analysed by existing technologies.

## Results

### Design of Tocky system for analysing the time and frequency domains of signal-triggered activation and differentiation events

Given the long half-life of stable fluorescent proteins (FP), like GFP (56 h (Sacchetti et al., 2001)), the dynamics of gene transcription cannot be effectively captured using conventional FP expression as a reporter. We therefore chose to use Fluorescent Timer protein (Timer), which forms a short-lived chromophore that emits blue fluorescence (Blue), before producing the mature chromophore that emits red fluorescence (Red). The maturation half-life (i.e. the production of red-form proteins) is estimated to be 7 h, while red-proteins are stable and have a decay rate longer than 20 h (Subach et al., 2009) (**Fig. 1A)**.

**Figure 1:**
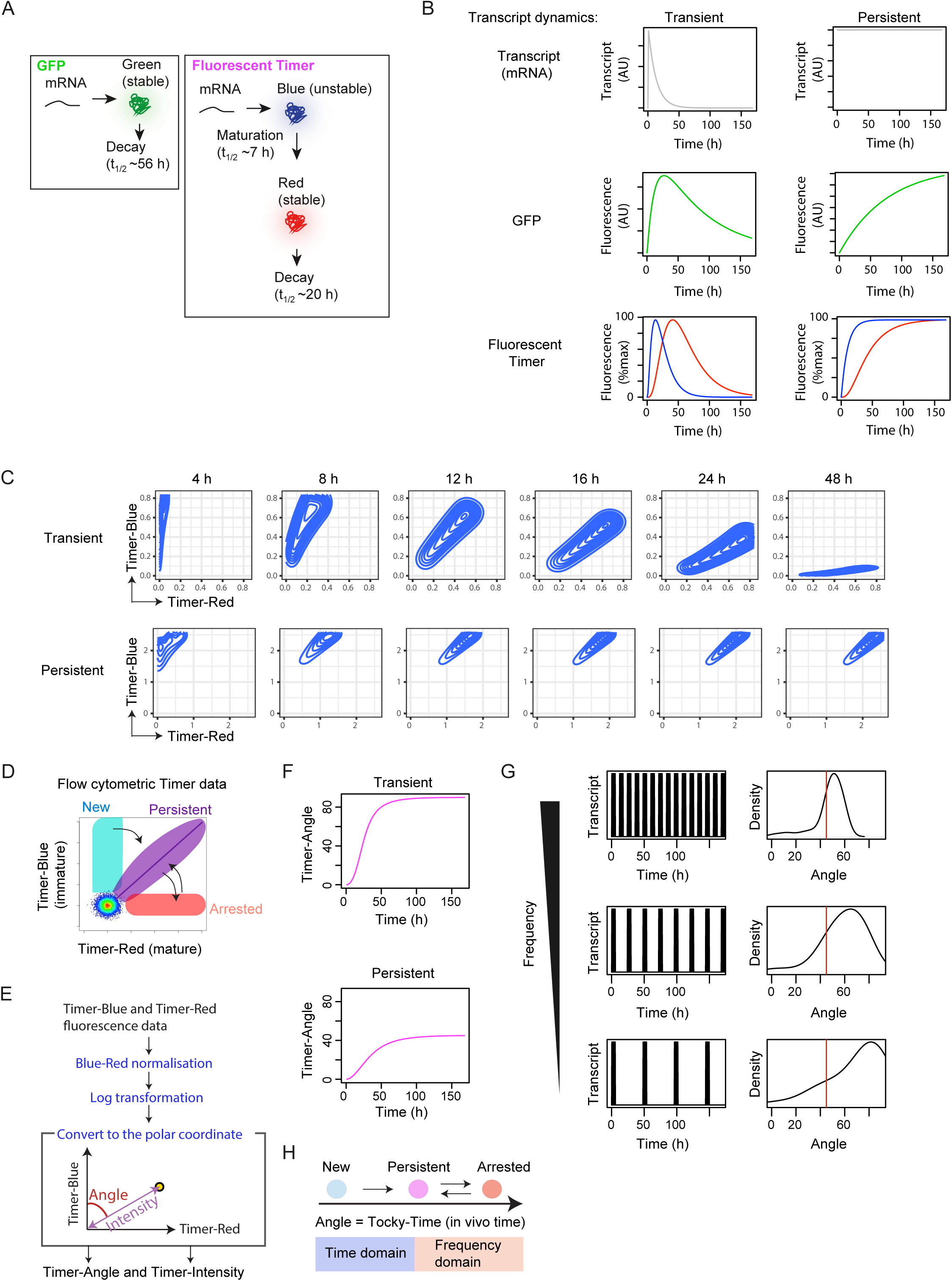
Design of Tocky system for analysing the time and frequency domains of signal-triggered activation and differentiation events. **(A)** Production and decay of GFP and Fluorescent Timer protein. **(B)** In silico analysis of GFP and Timer fluorescence by different transcriptional dynamics. Time course analysis of GFP or Blue and Red forms of Timer protein, simulated flow cytometric data, and Timer-Angle, given the constant influx of cells with the indicated transcriptional dynamics. Data from spike-like (transient) and constant transcriptional dynamics are shown. **(C)** In silico modelling of flow cytometric data depicting transient (top) or persistent Timer transcription (bottom). **(D)** Timer locus approach to translate flow cytometric Timer fluorescence data into transcriptional dynamics. **(E)** Schematic representation of Trigonometric data transformation of flow cytometric Timer data. Flow cytometric Blue and Red Timer fluorescence data were preprocessed and normalised, and subsequently transformed by a trigonometric function. **(F)** Effect of persistent or transient transcription on Timer Angle progression. **(G)** In silico modelling of the effect of transcriptional frequency on the distribution of Timer Angle values. **(H)** Model for analysing the time domain and frequency domain using Tocky-time analysis.

We investigated *in silico* how two different transcriptional activities would influence the production of blue and red fluorescent form of the protein through time, and how the light emitted would be detected by flow cytometry. We compared the reporting of transient, pulse-like transcription with that of persistent transcription, both of which are relevant to TCR-mediated transcription (Yosef and Regev, 2011) (**Fig. 1B)**. A linear kinetics model showed that Blue was a better readout for the real-time level of transcription than Red and GFP, which rather reported the cumulative activity of transcription. Next, we analysed the dynamics of Timer-expressing cells in the Blue-Red plane, which is relevant for flow cytometric analysis. Assuming that cells received a transient signal in a synchronised manner, cells showed a fan-like movement from Blue+Red- to Blue-Red+, and cells stayed until the red-form proteins decay. On the other hand, when cells receive persistent TCR signals, cells gradually approached the diagonal line between Blue and Red axes, which is the steady state (**Fig. 1C)**. Thus, there are 3 key loci in the Blue-Red plane (**Fig. 1D)**. Firstly, when new transcription occurs in Timer-negative cells, Timer^+^ cells acquire pure Blue, and are identified in the New locus. Secondly, if transcriptional activity is persistent and/or sufficiently frequent, cells accumulate in and around the steady-state diagonal line (Persistent locus). Lastly, when transcription is diminished, cells lose Blue and stay in the Arrested locus, until the red protein decays (**Fig. 1D)**. Importantly, Timer maturation is unidirectional and irreversible as the chromophore matures from Blue to Red. In the case of individual Timer^+^ cells, movement from New to Persistent loci is also unidirectional and irreversible, because the half-life of red protein is longer than the half-life of the blue form, whether the signal is transient or continuous (**Fig. 1C)**. Therefore transition from the New to the Persistent loci captures the time domain of cellular differentiation. In contrast, cells in the Arrested locus may re-initiate transcription to express new Blue protein and move anti-clockwise back into the Persistent locus (**Fig. 1D)**. This leads to the hypothesis that the movement between Persistent and Arrested loci more specifically captures how frequently transcriptional activities occur in mature cells.

These three loci can be identified and quantified more effectively by analysis of the angle of individual cells from the blue axis. The 2-dimensional blue vs. red Timer fluorescence data can be transformed by trigonometric data transformation and converted into the polar coordinate to provide new variables: the angle from the blue axis is defined as Timer-Angle, which is a measure of the trajectory and change in transcriptional history. The Euclid distance from the origin is defined as Timer-Intensity (**Fig. 1E)** and is a measure of the signal strength given Timer-Angle value.

In fact, trigonometric data transformation quantitatively showed that transient signals had a faster progression of Timer-Angle than continuous signals (**Fig. 1F)**. Next, using Timer-Angle, we analysed transcriptional activities with different frequencies. As implicated by the analysis of transient and continuous signals (**Fig. 1B** and **1C)**, when the frequency of transcription was low, cells accumulate in higher Timer-Angle. In contrast, if cells receive high-frequency signals, they approach the steady-state of continuous transcription (**Fig. 1G)**.

We therefore coined the system **T**imer **o**f **C**ell **K**inetics and Activit**y** (Tocky; 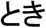 [toki] means “Time” in Japanese). Timer-Angle provides time-related composite information: the first phase (New ➔ Persistent) is for analysing how differentiation mechanisms are regulated over time (i.e. time domain analysis); and the latter half (Persistent – Arrested) is for analysing the relative frequency of transcriptional activation (i.e. frequency domain analysis). Thus, we designate this composite time-axis as *Tocky-Time* (**Fig. 1H**). In Time Domain analysis in physics and related subjects, data obtained over time are transformed into frequency data (and vice versa) using a function such as the Fourier transform. In contrast, the Tocky-Time can be the measurement of either time or frequency, depending on the Timer maturation phase. Notably, these two domains merge in the Tocky-time when transcriptional activities are persistent.

### The development of Nr4a3-Tocky for the analysis of TCR signal downstream events

T cell differentiation is triggered by TCR signals. In order to track the in vivo dynamics of transcription downstream of TCR signal transduction, we firstly identified genes immediately downstream of TCR signalling using a data-oriented multidimensional analysis, Canonical Correspondence Analysis (CCA) (Ono et al., 2014) (**Supplementary Fig. 1A)**. *Nr4a3 (Nor1)* was identified as the gene with the highest correlation with anti-CD3 mediated T cell activation (which mimics TCR activation) and in vivo TCR signals in the thymus, while *Nr4a1 (Nur77)* and *Rel* also scored highly (**Supplementary Fig. 1B** and **1C)**. In agreement, upon anti-CD3 stimulation, *Nr4a3/Nor1* was rapidly induced in T cells and peaked within 2 hours of stimulation (**Supplementary Fig. 1D)**. Having established *in silico* that Tocky can temporally report transcription, we generated BAC transgenic reporter *Nr4a3-* Tocky mice (**Fig. 2A)**, in which the transcriptional activity of the *Nr4a3* gene is reported by Timer proteins and used as an indicator of new transcription mediated by TCR signal transduction.

**Figure 2:**
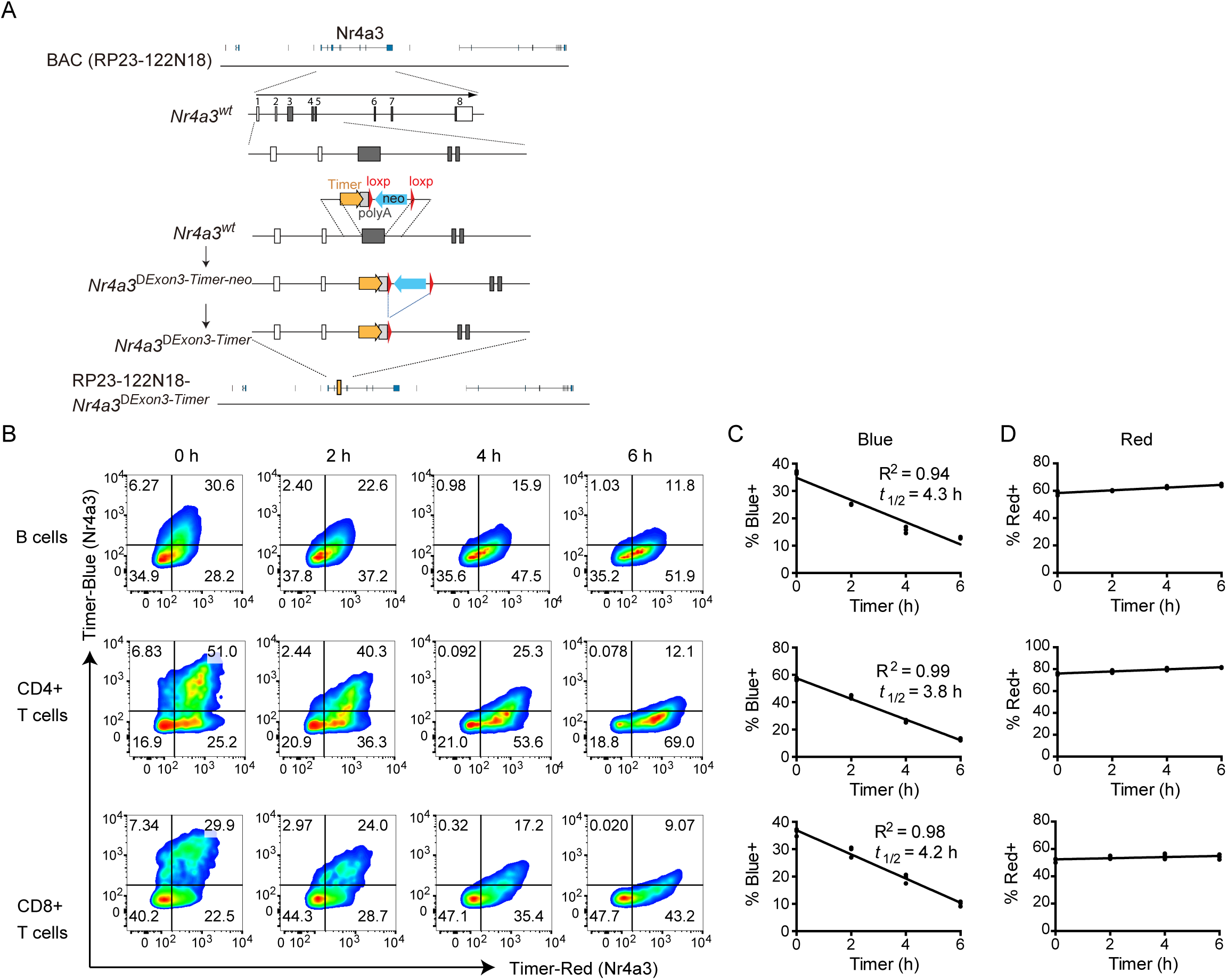
Antigen-receptor ligation induces Timer proteins in B and T-cell subsets from *Nr4a3*-Tocky mice. **(A)** Construct for generating Nr4a3-Tocky BAC transgenic mice. **(B)** Splenocytes from Nr4a3-Tocky mice were activated for 20 h with either 10μg/ml of soluble goat anti mouse IgM (for CD19+ B cells) or 2 μg/ml plate bound anti-CD3 (for CD4+ and CD8+ T-cells). Cells were then incubated with 100ug/ml cycloheximide to inhibit new protein translation and the decay of blue fluorescence measured over time by flow cytometry. Shown is Timer-Blue vs Timer-Red fluorescence in CD19+ B cells (top), CD4+ T-cells (middle) or CD8+ T-cells (bottom) at the indicated time points. **(C)** Summary data of the % cells Blue+ or **(D)** % Red+ in the cultures. Linear regression by pearson’s correlation. N=3 culture triplicates. Data are representative of two independent experiments.

### Antigen-receptor ligation induces Timer proteins in B and T-cell subsets from Nr4a3-Tocky mice

Using Nr4a3-Tocky, antigen-receptor ligation induced Timer protein expression not only in CD4+ T cells but also in CD8+ T cells and B cells, with the most prominent induction in CD4+ T cells (**Fig. 2B)**. This provided us an opportunity to experimentally determine the half-life of the blue fluorescent form of Timer protein in each cell subtype, and thereby ask if the kinetics of Blue-expressing cells are similar between different cell subtypes. B and T cells were stimulated by antigen-receptor ligation and immediately after the induction of Timer proteins, protein translation was blocked by cycloheximide and the decay of Blue fluorescence was measured by flow cytometry (**Fig. 2C)**. Linear regression analysis showed that the half-life of Blue-expressing cells was around 4 h in all the cell subtypes, while the percentage of Red-expressing cells was barely changed or slightly increased after the conversion of Blue proteins into Red proteins (**Fig. 2D)**. These data support that Timer maturation is not affected by cell subtypes.

### Nr4a3-Tocky reveals the temporal dynamics and frequency of TCR signal-triggered activation events

In order to analyse the T cell response upon antigen recognition, we generated OT-II *Nr4a3*-Tocky mice, which express ovalbumin (Ova)-specific transgenic TCR. Ova stimulation of CD4+ T-cells from OT-II *Nr4a3*-Tocky mice resulted in the upregulation of Blue within 4 h, with some cells acquiring Red at 8 h after stimulation (**Fig. 3A)**. *Nr4a3*-Tocky T cells further increased both Blue and Red throughout the 48-h culture, while removal of TCR stimulation by anti-MHC II-treatment from 24 h onwards resulted in the rapid loss of Blue (**Fig. 3A)**, which captured the reduction in frequency of TCR signalling.

**Figure 3:**
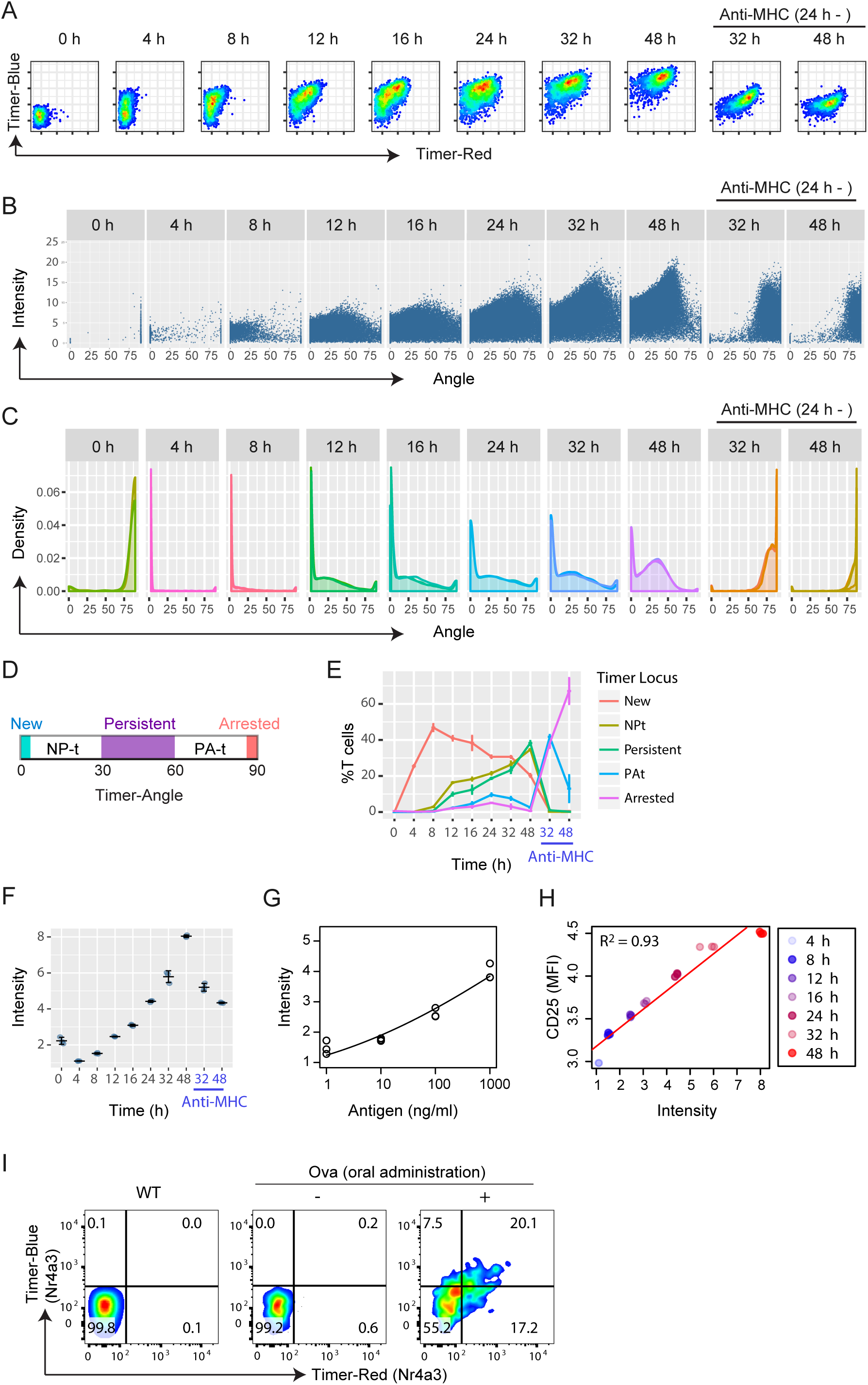
*Nr4a3*-Tocky reveals the time domain of TCR signal-triggered activation and differentiation events. **(A)** Flow cytometric analysis of Blue and Red Timer raw fluorescence in antigen-stimulated OT-II *Nr4a3*-Tocky T-cells. T cells were stimulated for the indicated time points with 1 μM Ova peptide. In some cultures, anti-MHC II antibodies were added at 24 h in order to terminate TCR signalling. **(B)** Trigonometric-transformed data of a. Individual cells are plotted against Timer-Angle and –Intensity. **(C)** Density plot of Timer-Angle from the transformed data. **(D)** The designation of the 5 Timer loci by Timer-Angle θ as follows: New (θ = 0°), NP-t (0° < θ < 30°), Persistent (30° ≤ θ < 60°), PA-t (60° ≤ θ < 90°) and Arrested (θ = 90°). **(E)** Timer Locus analysis to show the frequency of cells within the 5 Timer loci defined in **(C)**. **(F)** Summary of Timer-Intensity in the cultures from **(A)** over time. **(G)** Dose response curve of Timer-Intensity on stimulation with titrated doses of Ova peptide (Antigen). OT-II *Nr4a3*-Tocky T-cells were stimulated for 22 h in the presence of 1, 10, 100 or 1000 nM Ova peptide and APCs. Data were fitted to a dose-response curve with a statistical significance by a lack-of-fit test (p = 0.014). **(H)** Scatter plot of Timer-Intensity vs cell surface CD25 expression. Linear regression analysis showed a strong correlation (R^2^ = 0.93). See legend for sample identities. **(I)** CD4+ T-cells from OT-II TCR transgenic Nr4a3-Tocky mice were adoptively transferred in to CD45.1 congenic mice, which were fed ovalbumin or control for 3 days. CD45.2+CD45.1+ CD4+ OT-II *Nr4a3*-Tocky T-cells within mesenteric LN were analysed for Timer expression. Bars represent mean ± SD. N = 3 culture triplicates. Data are representative of at least two independent experiments.

In order to analyse effectively the continuous progression of Timer fluorescence in **Fig. 3A**, Timer fluorescence data were transformed into Angle and Intensity data (**Fig. 3B)**. This transformation of the data normalises for blue and red fluorescence by using the mean and standard deviation of the ‘negative cloud’ of cells (i.e. cells with autofluorescence only). In order to generate robust angle values, thresholding of the data is required to restrain angles between 0 and 90. This thresholding sets the blue and red levels for positivity, and thus collapses pure blue and pure red cells into angles of 0 and 90 respectively (See methods section).

Importantly, the progression of Timer Angle becomes slower and cells accumulate as cells approach ~45° by density plots, which visualise the distribution of angle values, reflecting sustained or high frequency TCR signalling (**Fig. 3C)**. In order to quantify the maturation of the Timer chromophore since first onset of its transcription, we grouped cells into 5 populations (i.e. data categorisation) according to their Angle values: from the ‘New’ population (Angle = 0°) representing cells within the first 4 hours of initiation of TCR-mediated transcription to the ‘Arrested’ population (Angle = 90°), which represents cells in which all Timer protein has matured to red, and transcriptional activity has decreased below cytometer detection thresholds (**Fig. 3D)**. The area between 30° – 60° was defined as the Persistent locus. The areas between New and Persistent (NP-t) or between the Persistent and Arrested (PA-t) contained cells from the surrounding loci that are in the process of changing their frequency of *Nr4a3* transcription (**Fig. 3E)**. The analysis of the percentage of cells in these loci (designated as Timer locus analysis) neatly captured the change in TCR-mediated transcription through time, as cells shifted from New to Persistent, which represent the time domain. Furthermore, it showed that removal of TCR signals lead to a complete loss of New, NP-t and Persistent signalling within 8 hours, and cells migrated to Arrested transcriptional dynamics, which represents sparse or no signalling activity (**Fig. 3E)**. Thus, *Nr4a3*-Tocky recaptured the predicted kinetics of Timer-expressing cells, validating the Tocky model (**Fig. 1H)**.

We hypothesised that Timer-Intensity reflects both the signal strength and the duration/ frequency of TCR signalling, as Timer proteins accumulate in individual cells in response to strong and/or repeated TCR signals. Timer Intensity in antigen-stimulated Nr4a3-Tocky T cells increased over time, and fell after removal of the TCR signal (**Fig. 3F)**. Furthermore, Timer Intensity was increased in a dose-dependent manner by cognate antigen (**Fig. 3G)**. Interestingly, Timer Intensity showed a high correlation to cell surface CD25 expression, which is a marker of activated T cells (R^2^ = 0.93) (**Fig. 3H)**. Thus, using Nr4a3-Tocky, Timer Intensity reflects the cumulative transcriptional outcome of signals in a given cell, as the reporter protein accumulates in response to sustained or highly frequent signals.

Next, we addressed whether Nr4a3-Tocky identifies T cells that receive TCR signals in vivo. We adoptively transferred OT-II TCR transgenic Nr4a3-Tocky T cells into congenic recipients, which were then fed their cognate antigen, Ova, in the water (or water alone) (**Fig. 3I)**. This model allows analysis of T cell responses in mesenteric lymph nodes to orally administered antigens. Timer expression occurred in OT-II TCR transgenic T-cells from Nr4a3-Tocky mice only in the presence of their cognate antigen, indicating that Timer expression is induced upon antigen recognition in vivo.

To validate the frequency domain of the Tocky system, we employed the Nr4a3-Tocky OTII system to perform periodic stimulation of T-cells with cognate peptide. T-cells underwent one, two or three rounds of 4 h peptide stimulation over a 48-h period (the groups I, II, and III), and the Timer Blue vs. Red expression was analysed by flow cytometry and compared to constant stimulation (Constant) (**Fig. 4A)**. As predicted, low frequency stimulation resulted in predominantly pure red expression (**Fig. 4B)**. With increased frequency of signaling, blue fluorescence increased, which was highest in cells undergoing constant stimulation. Analysis of Timer-Angle distribution captured the shift in frequency, as Angles moved from the Arrested locus (i.e. 90 degrees) towards the Persistent locus with increasing frequency of stimulus (**Fig. 4C** and **4D)**. As expected, Timer-Intensity also showed a frequency dependent relationship, as more Timer proteins accumulated in response to more frequent TCR stimulation (**Fig. 4E)**.

**Figure 4:**
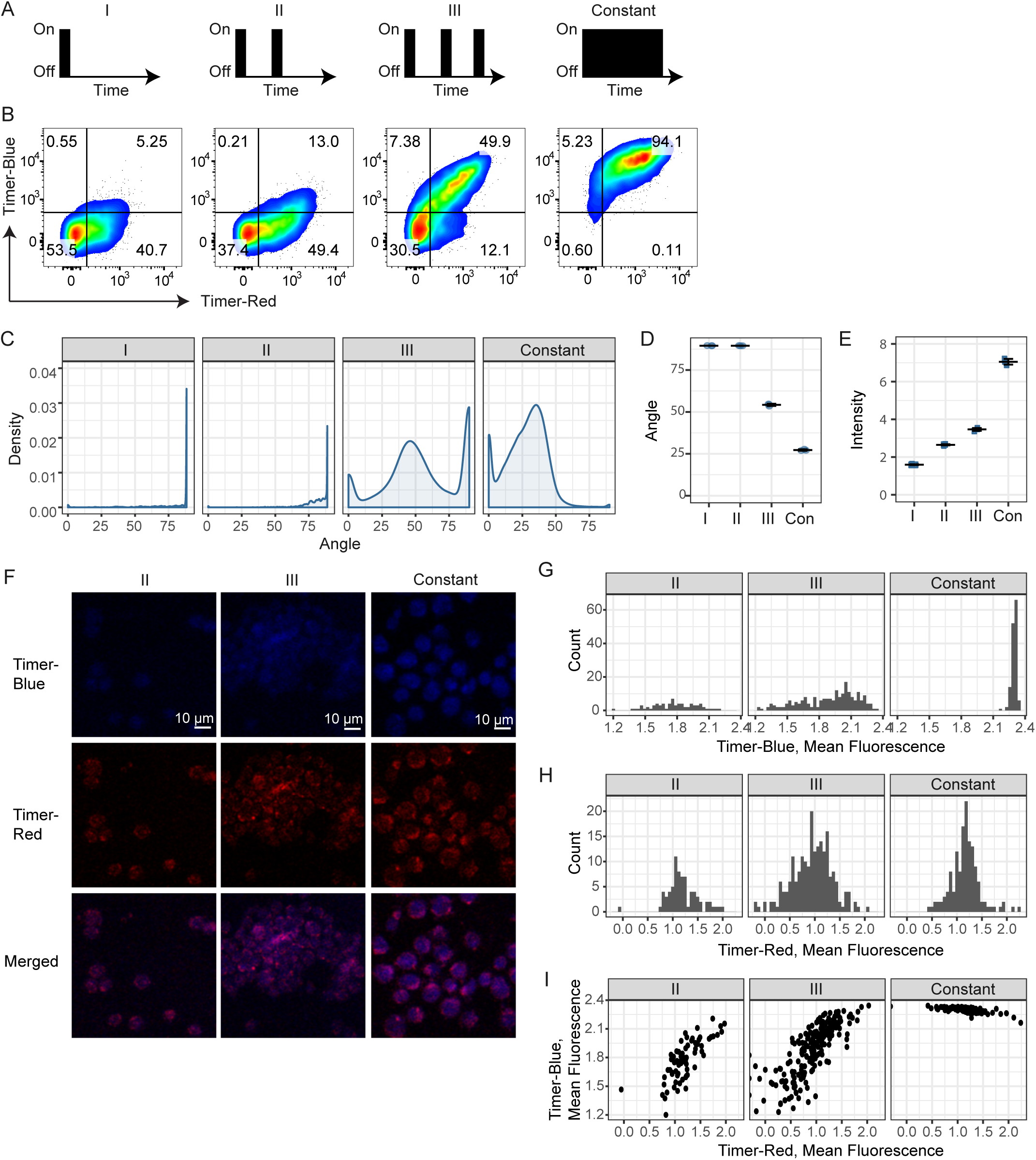
*Nr4a3*-Tocky captures the frequency domain of TCR signalling. T-cells from OTII *Nr4a3*-Tocky mice were cultured with T-cell depleted splenocytes. **(A)** T-cells were stimulated with 1 μM ova for either **(I)** 4 h then rested for 48 h; (II) 4 h stimulation, 20 h rest, 4 h stimulation, 24 h rest (III) 4 h stimulation, 20 h rest, 4 h stimulation, 20 h rest, 4 h stimulation or (Con) 2 days constant stimulation. During resting stages, cells were incubated with 100U/ml recombinant IL-2 and 20 μg/ml anti-MHC II, and washed before and after stimulations. **(B)** Cells were then harvested and CD4+ OTII T-cells analysed for Timer-Blue vs. Timer-Red expression by flow cytometry. **(C)** Displayed are the Timer-Angle distributions for the four culture conditions. Shown are the mean **(D)** Timer-Angle or Timer-Intensity **(E)** within the 4 different cultures. Bars represent mean ± SD. N = 3 culture triplicates. **(F)** Confocal microscopy analysis of OTII *Nr4a3*-Tocky stimulated with various frequencies. The same cell samples used in A-D were analysed by confocal microscopy. Bars indicate 10 μm. (G - H) Histogram showing the mean fluorescence intensity of **(G)** Timer-Blue or **(H)** Timer-Red in single cells by microscopic image analysis. **(I)** Two dimensional plots of the mean fluorescence intensities of Timer-Blue and Timer-Red in single cells by microscopic image analysis. Cells were identified as Region of Interest (ROI) and mean fluorescence intensities were measured and logged. See **Supplementary Fig. 2** and Material and Methods for the details.

Next, we have further analysed the same stimulated T cells using confocal microscopy analysis (**Fig. 4F-4I**, **Supplementary Fig. 2A** and **2B)**. Visual inspections of microscopic images suggested that Timer-blue fluorescence was increased as the frequency of stimulation increased (**Fig. 4F**, **Supplementary Fig. 2A)**. Quantitative measurement of fluorescent intensities in individual single cells confirmed this (**Supplementary Fig. 2B** and **Fig. 4G – 4I)**. Both Timer-blue and -red fluorescence increased as the stimulation was more frequently applied, and the increase was more remarkable in Timer-blue fluorescence (**Fig. 4G, 4H)**. Two-dimensional plot of Timer-blue and –red fluorescence by confocal microscopy (**Fig. 4I)** recaptured the flow cytometric result (**Fig. 4B)**. Thus, this single cell microscopy experiment further validated the Tocky system.

### Cell division, co-stimulation, and IL-2 signalling do not affect Timer-Angle progression

Next, we examined whether processes related to T cell activation affect Timer-Angle. TCR signalling leads to T cell activation and proliferation. Since each cell division halves both existing blue- and red-form proteins, it was predicted that cell division will not change Timer-Angle. In fact, by analysing the dilution of a proliferation dye as cells divide, activated T cells did not change their Timer-Angle after cell division (**Fig. 5A** and **5B**). Activated T cells produce IL-2, which promotes the survival and proliferation of these T cells (Hoyer et al., 2008). Exogenous IL-2 had no effect on Timer-Angle (**Fig. 5C** and **5D**). CD28 signalling enhances the activities of TCR signal downstream, inhibiting apoptosis and sustaining the activation processes (Boomer and Green, 2010; Walker and Sansom, 2011). Importantly, anti-CD28 antibody alone did not induce Timer expression, and also, it did not change the progression of Timer-Angle by TCR signals (**Fig. 5E** and **5F)**.

**Figure 5:**
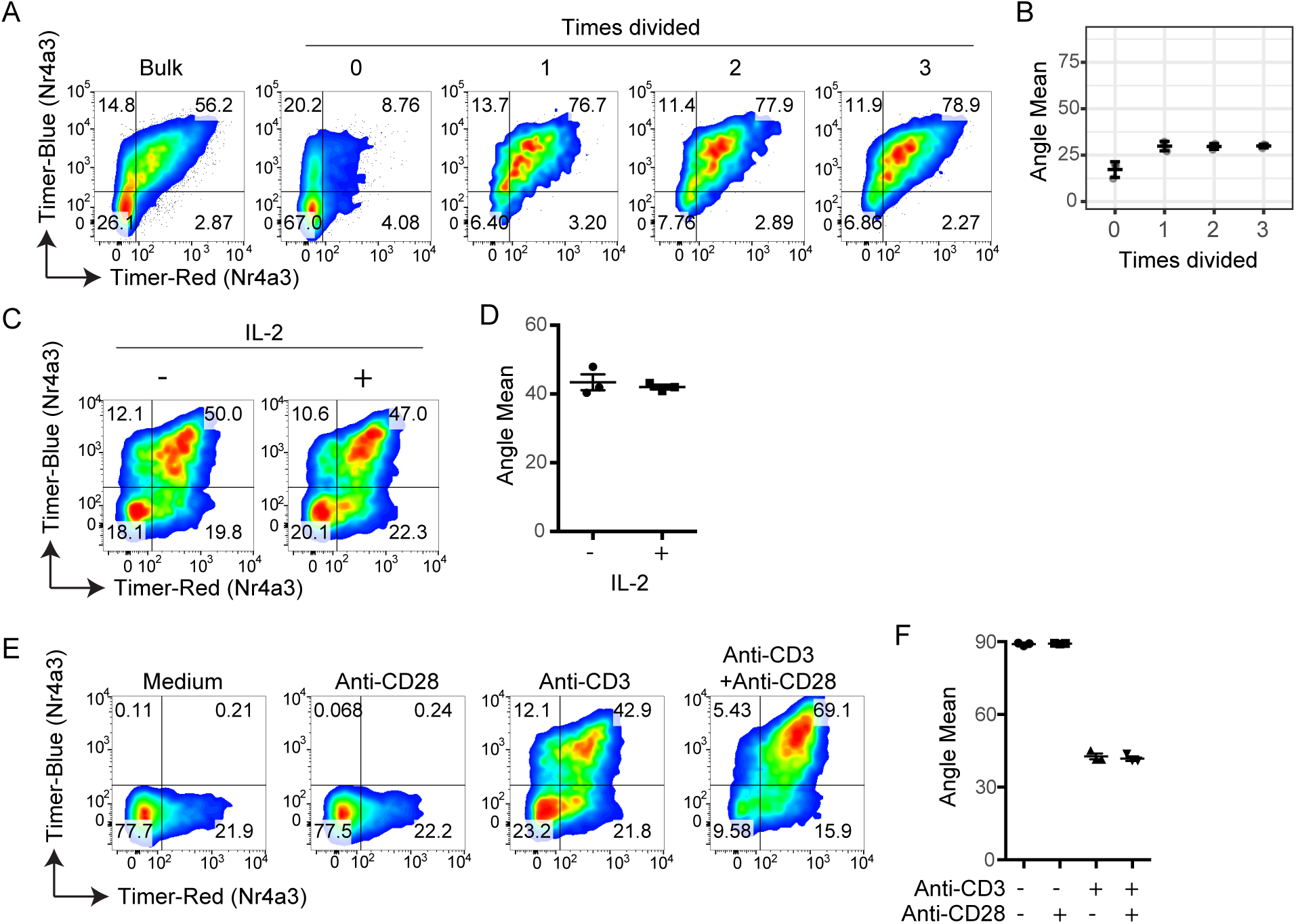
Cell division, co-stimulation, and IL-2 signalling do not affect Timer-Angle progression. CD4+ T-cells from *Nr4a3*-Tocky mice were labelled with a proliferation dye and activated for 72 h with anti-CD3. Cells were then analysed based on dilution of proliferation dye, and classified into number of cellular divisions. **(A)** Shown is Timer-Blue vs Timer-Red fluorescence in CD4+ T-cells gated on dilution of proliferation dye. **(B)** Mean Timer Angle values in the cultures from **(A)**. (C-D) Splenocytes from *Nr4a3*-Tocky mice were stimulated on anti-CD3 coated plates in the presence or absence of 100U/ml rhIL-2 for 20 h. **(C)** Shown are Timer-Blue vs. Timer-Red fluorescence in CD4+ T-cells from cultures. **(D)** Mean Timer Angle in cultures. N=3 culture triplicates, bars represent mean+/-SEM. Data are representative of two independent experiments. (E-F) Splenocytes from *Nr4a3*-Tocky mice were stimulated on plates coated with anti-CD28 alone, anti-CD3 alone or anti-CD3+anti-CD28 for 20 h. **(E)** Shown are Timer-Blue vs. Timer-Red fluorescence in CD4+ T-cells from cultures. **(F)** Mean Timer Angle in cultures. N=3 culture triplicates, bars represent mean+/-SEM. Data are representative of two independent experiments.

Collectively, the data above indicate that the progression of Timer-Angle is not affected by cell division or activation status, but defined by the time and signal dynamics, and that Nr4a3-Tocky reports the temporal dynamics of TCR signal downstream activities.

### The time-domain analysis of Nr4a3-Tocky mice delineates the temporal sequences of thymic Treg differentiation in vivo

Having validated the *Nr4a3*-Tocky system, we decided to investigate thymic Treg differentiation. TCR signaling is the major determinant of regulatory T-cell (Treg) differentiation in the thymus. T cells that have recognised their cognate antigens and received strong TCR signals preferentially express CD25 and Foxp3 and differentiate into Treg (Hsieh et al., 2012; Weissler and Caton, 2014).

In order to demonstrate the quantitative power of the Tocky technology, we investigated the temporal sequence of thymic CD4 T cell differentiation following TCR signals by analysing the time domain of *ex vivo* T cells from Nr4a3-Tocky thymus. Timer expression occurred in the CD4+CD8+ double positive (DP), CD4 single positive (SP), and CD8SP populations, with CD4SP displaying the highest frequency (8.9 ± 3.7%, **Fig. 6A)**. The cell surface expression of CD69, which is highly expressed on immature CD4SP, was high in CD4SP cells in the New locus, and progressively declined as the Timer protein matures. In contrast, the majority of CD4SP cells in the Persistent locus expressed Foxp3 and CD25 (**Fig. 6B)**. With the assumption that in neonatal mice, nearly all cells in the Persistent locus would be derived from the New locus, these results led to the hypothesis that Treg differentiation requires persistent TCR stimulation.

**Figure 6:**
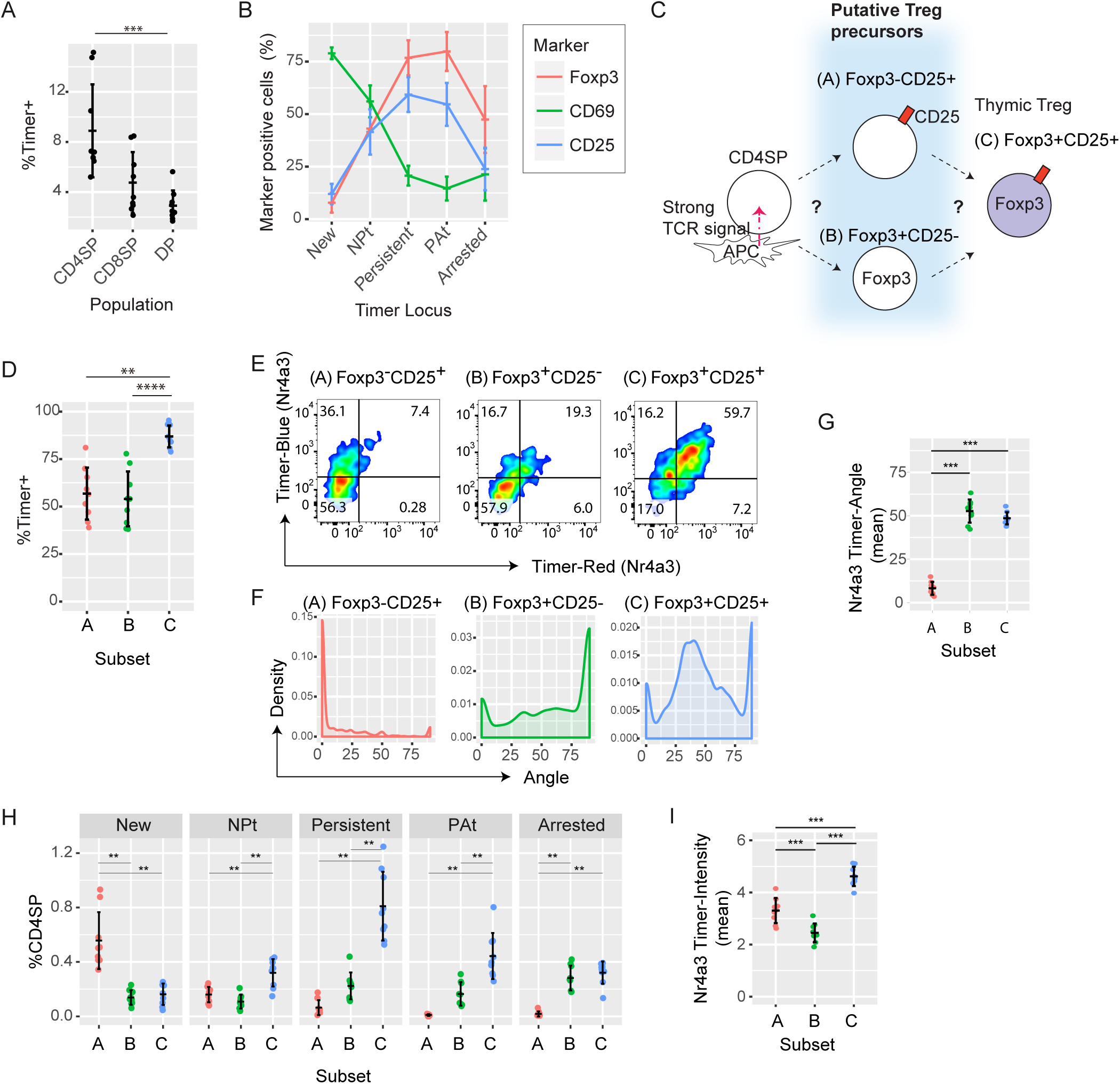
*Nr4a3*-Tocky mice identify thymic Treg precursors. **(A)** Percentages of Timer^+^ cells in the indicated thymic T cell populations from *Nr4a3*-Tocky mice. **(B)** Mean percentages of CD4SP cells expressing CD69 (green), CD25 (blue) or Foxp3 (red) from the 5 Timer loci. **(C)** Current working model for thymic Treg differentiation. **(D)** Percentages of Timer^+^ cells in the indicated CD4SP subpopulations (A = CD25^+^Foxp3^-^, B = CD25^-^Foxp3^+^, C = CD25^+^Foxp3^+^). **(E)** Timer Blue and Red fluorescence from CD4SP Thymic Treg subsets from 7 day-old neonates. **(F)** Density plot of Timer-Angle from the transformed data. **(G)** The mean Timer-Angle of thymic Treg subsets. **(H)** Frequency within the CD4SP thymic population of each Timer locus in the three Treg subsets. **(I)** The mean Timer-Intensity of thymic Treg subsets. Thymi from 7-, 9- and 13-day old neonates were analysed in three independent experiments and data were combined, unless otherwise indicated. Bars represent mean ± SD.

The sequence of thymic Treg development is controversial. Some studies suggest that CD25^+^Foxp3^-^ Treg are the major Treg precursors (Burchill et al., 2007; Lio and Hsieh, 2008) while other studies argue that CD25^-^Foxp3^+^ cells are Treg precursors (Tai et al., 2013) (**Fig. 6C)**. These studies used in vitro culture experiments and intrathymic injection of precursor populations, which may not reflect the differentiation dynamics in vivo. We therefore revisited this issue to reveal the temporal sequences of thymic Treg differentiation in otherwise unmanipulated animals by analysing the time domain of Nr4a3-Tocky.

The majority of Treg (CD25^+^Foxp3^+^) were Timer^+^ (~90%), and both of the proposed Treg precursor populations (CD25^+^Foxp3^-^ and CD25^-^Foxp3^+^) also had high proportions of Timer^+^ cells (~63% and ~58%, respectively, **Fig. 6D)**, indicating that these three populations have all received either strong or frequent TCR signals. To place the three populations in time following TCR signal transduction, we analysed thymi from *Nr4a3-*Tocky neonates, and measured Foxp3 and CD25 expression, in addition to Timer fluorescence. Timer expressing CD25^+^Foxp3^-^ cells were mostly Blue+Red-, while those of CD25^+^Foxp3^+^ Treg and CD25^-^Foxp3^+^ cells were mostly Blue+Red+ (**Fig. 6E)**. Most of CD25^+^Foxp3^-^ cells had the Angle value 0, while CD25^+^Foxp3^+^ Treg showed a clear peak around 30–50° and CD25^-^Foxp3^+^ cells had a higher peak at 90° (**Fig. 6F)**. The mean of Angle was not significantly different between CD25^+^Foxp3^+^ Treg and CD25^-^Foxp3^+^ cells (**Fig. 6G)**. These indicate that the CD25^+^Foxp3^-^ population are in the earliest time after receiving TCR signals. Next, in order to define the temporal relationships between the three Treg and precursor populations, we quantified the proportion of Timer^+^ cells in each maturation stage of Timer fluorescence (**Fig. 6H)**. This showed that most of the CD4SP cells in the New locus were CD25^+^Foxp3^-^ cells, while most of CD4SP cells in the Persistent were CD25^+^Foxp3^+^ Treg. The relative contribution of CD25^-^Foxp3^+^ was greatest in the Arrested locus, suggesting that these cells are enriched with those with aborted TCR signalling (**Fig. 6H)**. In addition, Timer intensity analysis showed that the CD25^-^Foxp3^+^ subset had received the weakest and/or least frequent TCR signals among these subsets (**Fig. 6I)**.

Collectively, these analyses demonstrate that the major pathway for Treg differentiation is from CD25^+^Foxp3^-^ Treg precursors, in which persistent TCR signals induce Foxp3 expression. The CD25^-^Foxp3^+^ subset is enriched with Foxp3^+^ cells that have received relatively weaker and/or less sustained TCR signals. Following TCR signalling, cell surface CD69 expression peaks within 4 to 8 hours. CD25 expression is induced in this early phase in the Timer Blue (New) population and steadily accumulates as T cells receive TCR signals. Foxp3 expression is the most delayed and occurs most efficiently after T cells have persistently interacted with antigen (Fig. 7). Importantly, the Tocky system has for the first time directly shown the temporal sequences of Treg differentiation processes, revealing that temporally persistent TCR signals induce CD25^+^Foxp3^+^ Treg differentiation. Thus, our investigations show that Nr4a3-Tocky and Timer locus analysis effectively unravel the temporal dynamics of T cell differentiation following TCR signals.

**Figure 7.**
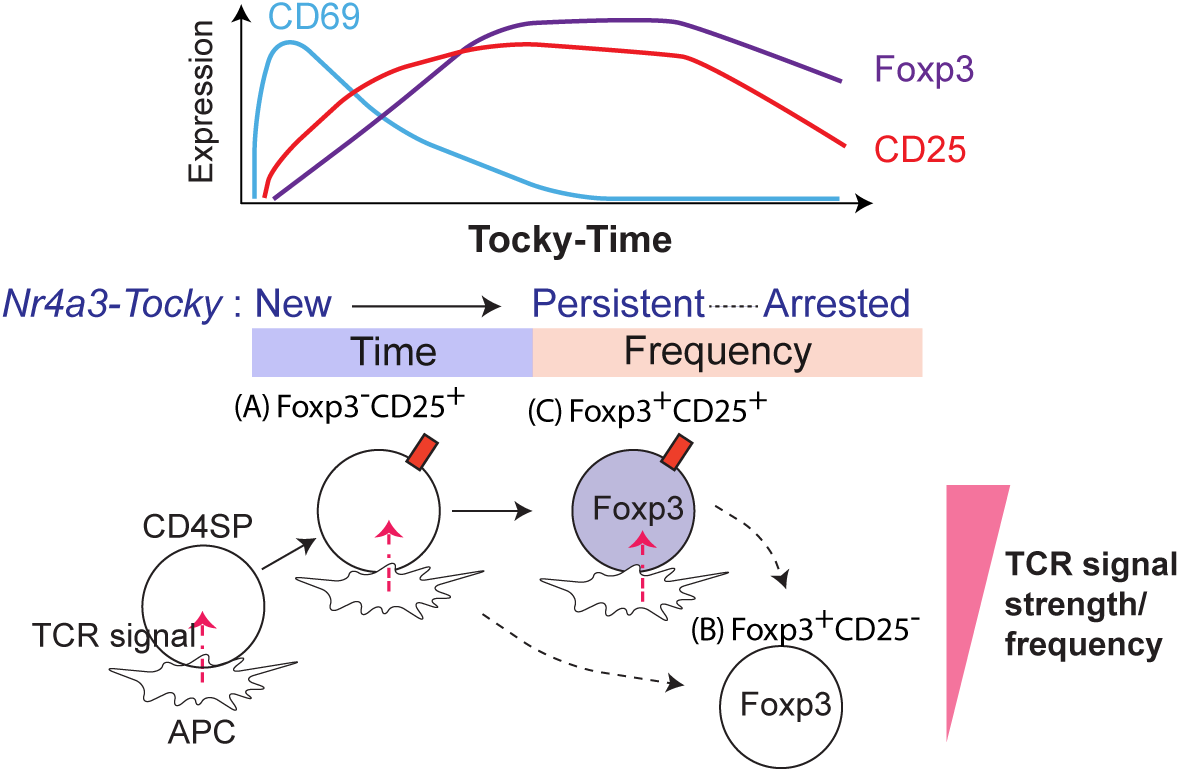
Model for the influence of TCR signaling dynamics on thymic Treg differentiation.

### Spontaneous and infrequent TCR signals occur in a minority of peripheral self-reactive T cells from Nr4a3-Tocky mice

Considering that Treg differentiation occurs through the recognition of cognate antigens in the thymus (Picca et al., 2006) and that these cells receive persistent TCR signals (Fig. 6), we hypothesised that Nr4a3-Tocky allows to identify by the persistent dynamics of TCR signals antigen-reactive T cells that recognise their cognate antigen during inflammation.

Firstly, we analysed *Nr4a3*-Tocky mice with WT polyclonal T cell repertoire, and found that Timer expression spontaneously occurred in a minority of the T cells from *Nr4a3*-Tocky mice and was mainly Blue-Red+ (**Supplementary Fig. 3A)**. Remarkably, Timer was expressed by a majority of Treg and memory-phenotype T cells (66.1 ± 5.1% and 30.8 ± 6.1%, respectively), while it was expressed by only a small proportion of naïve T cells (5.7 ± 2.5%) (**Supplementary Fig. 3B)**. Since both Treg and memory-phenotype T cells are self-reactive T cells (Ono and Tanaka, 2016), we hypothesized that all these Timer^+^ cells are in fact self-reactive T cells that spontaneously recognise self-antigens and receive TCR signals in the periphery. Interestingly, the Timer-Angle of most of Timer^+^ cells was 90° (the Arrested locus) irrespective of which T cell fraction they are from (**Supplementary Fig. 3C** and **3D)**. This indicates that all these Timer^+^ populations received infrequent TCR signals with short durations in a similar manner. In order to further confirm this, *Nr4a3*-Tocky T cells were adoptively transferred into congenic MHC Class II KO mice, to assess the contribution of spontaneous interactions between TCR and self-antigen/MHC Class II to Timer expression (**Supplementary Fig. 3E)**. As expected, 9 days after transfer, most of Timer expression was lost within MHC Class II KO mice. Collectively, these indicate that the spontaneous Timer expression in CD4+ T cells is induced through the infrequent and not-immunogenic interaction of self-reactive TCRs and self-antigen/MHC, which is currently called ‘tonic TCR signals’ (Klein et al., 2014; Ono and Tanaka, 2016).

### The frequency-domain analysis of Nr4a3-Tocky mice identify tissue-infiltrating antigen-reactive T cells as cells receiving persistent TCR signals

Next, we asked if immunogenic T cell responses have distinct dynamics of TCR signals compared to the infrequent TCR signals in self-reactive T cells. We used a murine model of Multiple Sclerosis, Experimental Autoimmune Encephalomyelitis (EAE), which produces autoimmune T cell responses to myelin basic proteins in the central nervous system (CNS) (O'Neill et al., 2006). Upon immunization of myelin oligodendrocyte glycoprotein (MOG), mice developed paralysis within 2 weeks, when T cells in the draining lymph nodes (dLN) of the immunized site and the spinal cord (CNS) were isolated and analysed (**Fig. 8A)**. MHC Class II tetramer for MOG-specific T cells (designated as MOG-tetramer) stained more than 10% of CNS-infiltrating T cells, while it stained only 0.5% of T cells in dLN (**Fig. 8B** and **8C)**. This indicates that most of cells in dLN cells are not reactive to MOG, while CNS-infiltrating T cells are markedly enriched with MOG-specific T cells, which mediate pathological inflammation (Stromnes and Goverman, 2006). Strikingly, almost 100% of CNS-infiltrating MOG-tetramer^+^ cells expressed Timer, while 40% of MOG-tetramer^+^ cells were Timer^+^ in dLN (**Fig. 8D)**. These findings show that a significantly greater proportion of CNS-infiltrating MOG-specific T-cells are actively engaged in TCR signals compared to the dLN. Next, we analysed the Timer expression in these cells. T cells in dLN, whether MOG-tetramer-positive or negative, showed prominent peaks at the New and Arrested loci (**Fig. 8E** and **8F)**. This suggests that a significant proportion of T cells receive TCR signals at each moment (therefore producing a peak at the New locus), while TCR signals in individual T cells are infrequent (hence producing a peak at the Arrested locus, c.f. **Fig. 1E)**. In contrast, CNS-infiltrating MOG-specific T cells were almost exclusively Blue+Red+, and had a single peak between 30–60 degree, while CNS-infiltrating tetramer-T cells had peaks in the New and Arrested loci (**Fig. 8E** and **8F)**, showing a similar pattern to those of dLN cells (**Fig. 8F)**. Most of the MOG-specific T cells in the CNS were found in the Persistent and NP-t loci (**Fig. 8G)**, further confirming that CNS-infiltrating MOG-specific T cells frequently receive TCR signals.

**Figure 8.**
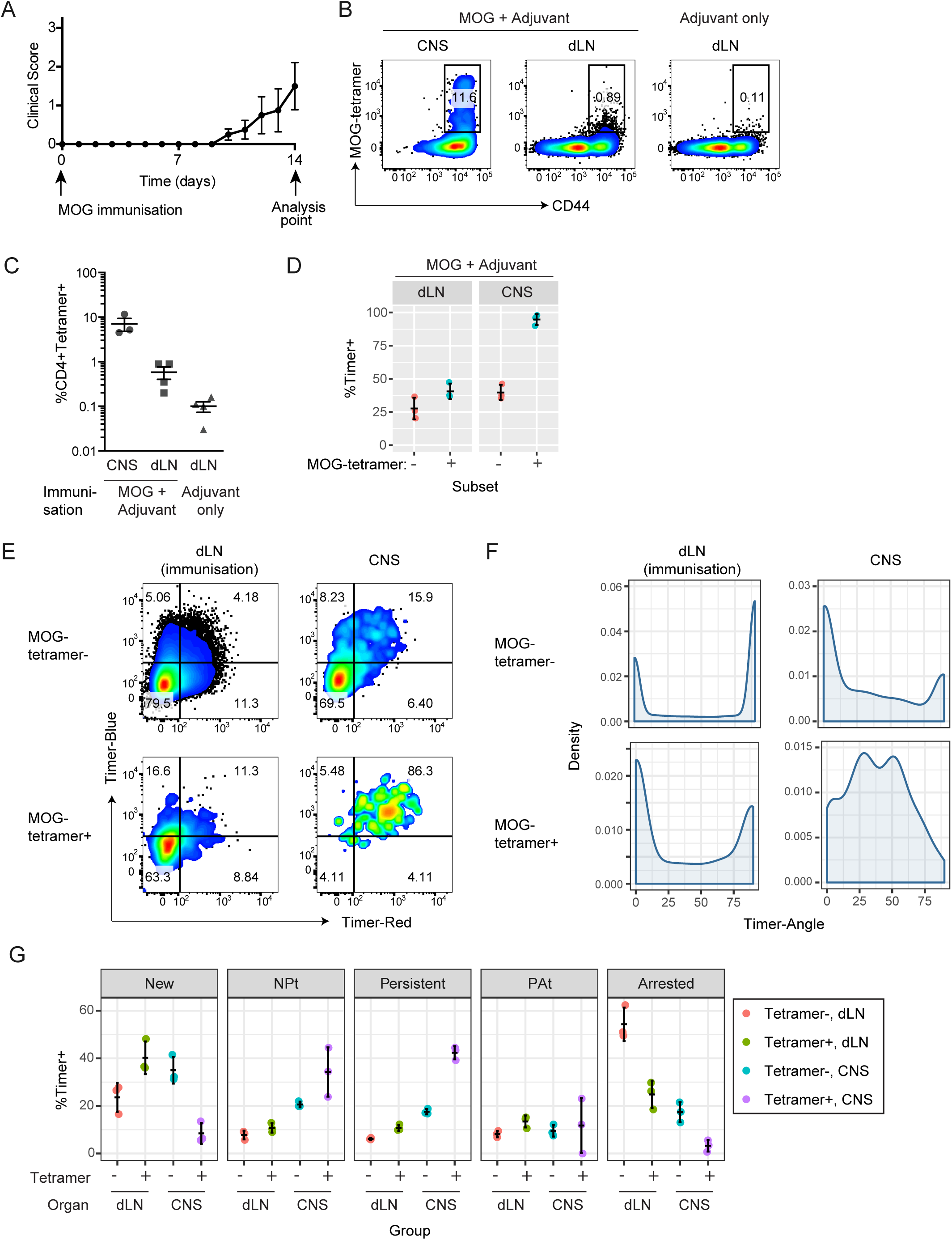
Tocky mice identify antigen-reactive T cells as cells receiving frequent TCR signals. **(A)** Development of clinical symptoms by MOG immunization. **(B)** At Day 14, CD4+ T-cells within the CNS or dLN were analysed for MOG-Tetramer labeling. Flow cytometry plots depict CD44 vs. MOG-Tetramer gated on CD4+ T-cells. **(C)** Percentages of MOG-tetramer^+^ cells in dLN of immunization site and the CNS from MOG immunized mice or dLN from control mice with adjuvant only. **(D)** Percentages of Timer^+^ cells in MOG-tetramer + and - cells in either dLN or CNS from MOG-immunised mice. (E-G) MOG-tetramer + and - cells in either dLN or CNS from MOG-immunised mice were analysed for **(E)** Blue and Red Timer raw fluorescence by flow cytometry, **(F)** Timer-Angle by density plot. **(G)** Timer Locus analysis of data from **(E)**. N=3 mice, bars represent mean+/-SD. Data are representative of two independent experiments.

Collectively, the results above indicate that MOG-specific T cells are engaged with their cognate antigens in the CNS, receiving frequent TCR signals. The accumulation of Blue+Red+ cells does not occur in LN, presumably because both MOG-specific T cells and MOG-presenting APCs are rare in LN. Thus, *Nr4a3*-Tocky identifies antigen-reactive T-cells by their transcriptional response in the nucleus. The signal dynamics of these antigen-reactive T cells are distinct from those of bystander or non-reactive T cells, and are intriguingly similar to those of thymic CD25^+^Foxp3^+^ Treg (**Fig. 6E versus 8E)**, which are engaged with self-antigen-presenting thymic APCs (namely, under agonistic selection (Klein et al., 2014)).

### *Foxp3-*Tocky *successfully identifies newly generated Treg*

Next, in order to address whether the Tocky system can be applied to another gene, and to further validate the system, we developed *Foxp3*-Tocky mice using the same approach as *Nr4a3*-Tocky (**Fig. 9A** and **9B)**. By investigating Foxp3 protein staining (**Fig. 9C)** and *Foxp3^IRES-GFP^ Foxp3-Tocky* double transgenic mice (**Fig. 9D)**, Timer expression showed high correlation with GFP and with Foxp3 protein. As expected, when naïve CD4^+^ T cells were stimulated in the presence of IL-2 and TGF-β (i.e. induced-Treg [iTreg] conditions), new Foxp3 expression (Blue^+^Red^-^) was induced at 22 h, which gained Red proteins by 50 h as the Timer chromophore matured (**Fig. 9E)**. Trigonometric Timer data analysis showed *ex-vivo* splenic Foxp3^+^ T cells from adult mice had high Timer-Angle values throughout the culture. In contrast, iTreg showed very low Angle values at 22 h, which slowly increased overtime (**Fig. 9F**–**9G)**. These data indicate that *Foxp3*-Tocky mice can identify the newly differentiating Treg that have recently initiated *Foxp3* transcription in vivo, which cannot be achieved through use of existing methods, such as GFP reporter mice.

**Figure 9.**
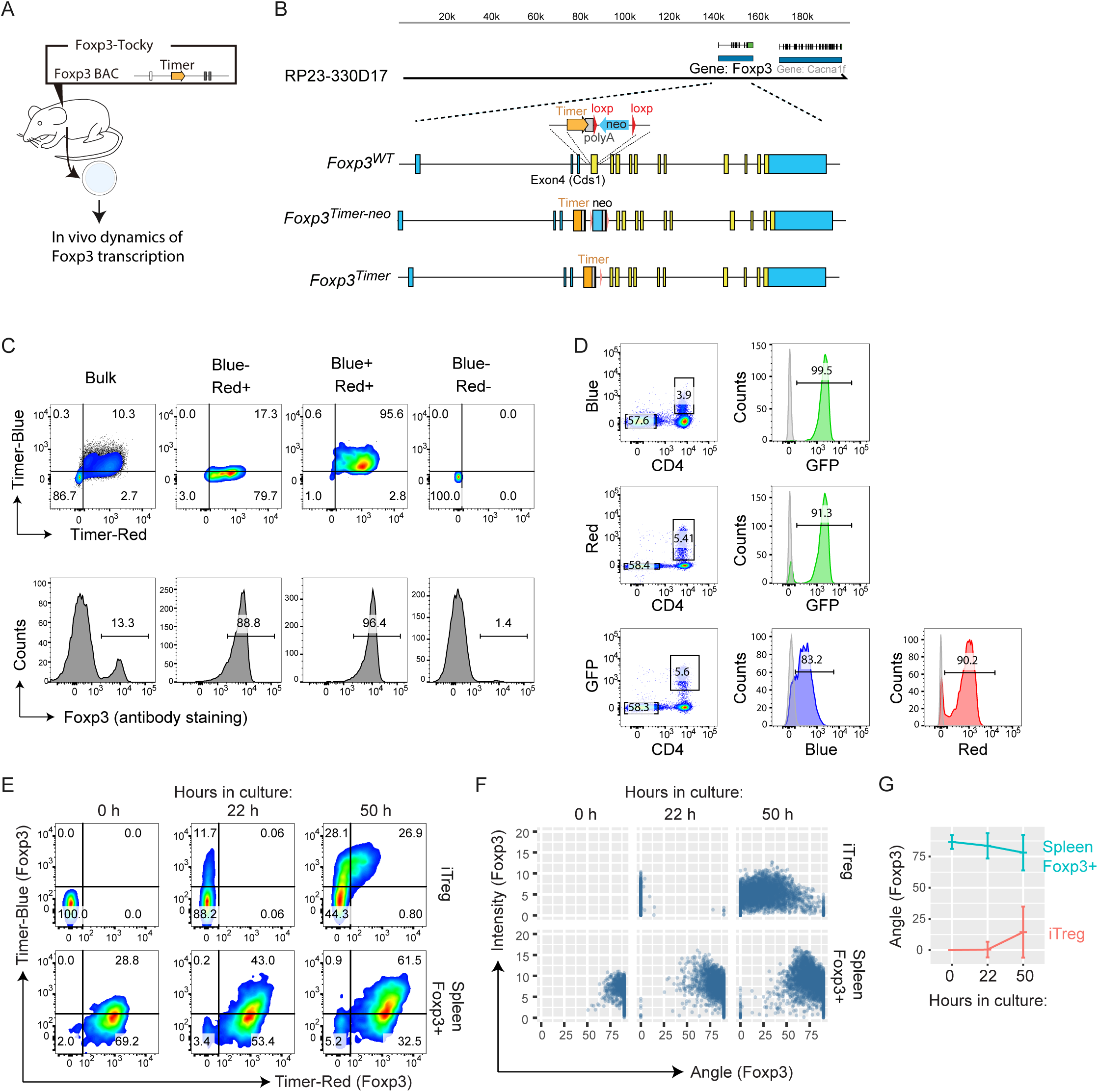
*Foxp3-Tocky* identifies newly generated Treg. **(A)** Use of *Foxp3*-Tocky mice to investigate in vivo dynamics of *Foxp3* transcription. **(B)** Construct for the generation of Foxp3-Tocky BAC transgenic mice. **(C)** Splenic T cells were sored into Blue-Red+, Blue+Red+, and Blue- Red- and analysed for intracellular Foxp3 proteins. **(D)** *Foxp3-Tocky* mice were crossed with Foxp3-IRES-GFP, and analysed for the co-expression of GFP and Timer. **(E)** Timer^−^ naïve T-cells and splenic Timer^+^ Treg from *Foxp3-* Tocky mice were isolated and stimulated by anti-CD3 and -CD28 for 0, 22 or 50 h in the presence of IL-2 (and TGF-β for iTreg). Flow cytometry plots display raw Blue vs Red expression during the cultures. **(F)** Timer Angle vs. Intensity, or **(G)** Timer Angle vs. Time in the data from E.

### Foxp3-Tocky reveals that the demethylation of the Foxp3 gene actively occurs when Foxp3 transcription is highly sustained in vivo

Next, taking advantage of the Tocky system, we investigated the transcription and methylation of the *Foxp3* gene during Treg generation. Whilst it is known that demethylation can occur within a few hours in cultured cells (Metivier et al., 2008), there is no available method to investigate the in vivo dynamics of demethylation in the mouse body. Previous work by the Huehn group have shown that immature Treg, as identified by high CD24 expression (Marodon and Rocha, 1994), display methylation of the Treg-Specific Demethylated Region (TSDR) of the *Foxp3* gene, and loss of CD24 expression is associated with its demethylation (Toker et al., 2013). However, since the in vivo dynamics of CD24 expression is not fully known, it is still unclear how the demethylation of the TSDR dynamically occurs in vivo. Thus, by analysing the time domain of *Foxp3* transcription using Foxp3-Tocky, we investigated the temporal dynamics of the demethylation of the TSDR during thymic Treg development. We isolated differentiating thymic T cells by flow-sorting Blue^high^ T-cells into Timer-Angle Low, Medium, and High (**Fig. 10A)**. Analysis showed that the isolated populations exhibited distinct Timer Angles from a mean of 10 to 80 degrees (**Fig. 10B & C**). DNA was isolated from each population and their TSDR demethylation was analysed (**Fig. 10D)**. Importantly, the newest Foxp3^+^ cells (Low) were still mostly methylated and not significantly different from Timer(-) cells in the degree of TSDR demethylation, while rapid demethylation occurred as cells moved to the Persistent locus (**Fig. 10D)**. Timer-Angles showed a strong correlation with TSDR demethylation rates (Spearman’s correlation coefficient ρ = -0.81, **Fig. 10E)**, indicating that the *Foxp3*-Tocky reporter successfully captures the demethylation dynamics of the *Foxp3* gene. Thus, Foxp3 expression precedes the demethylation of the TSDR region, and the most active demethylation process occurs when *Foxp3* transcription is sustained.

**Figure 10.**
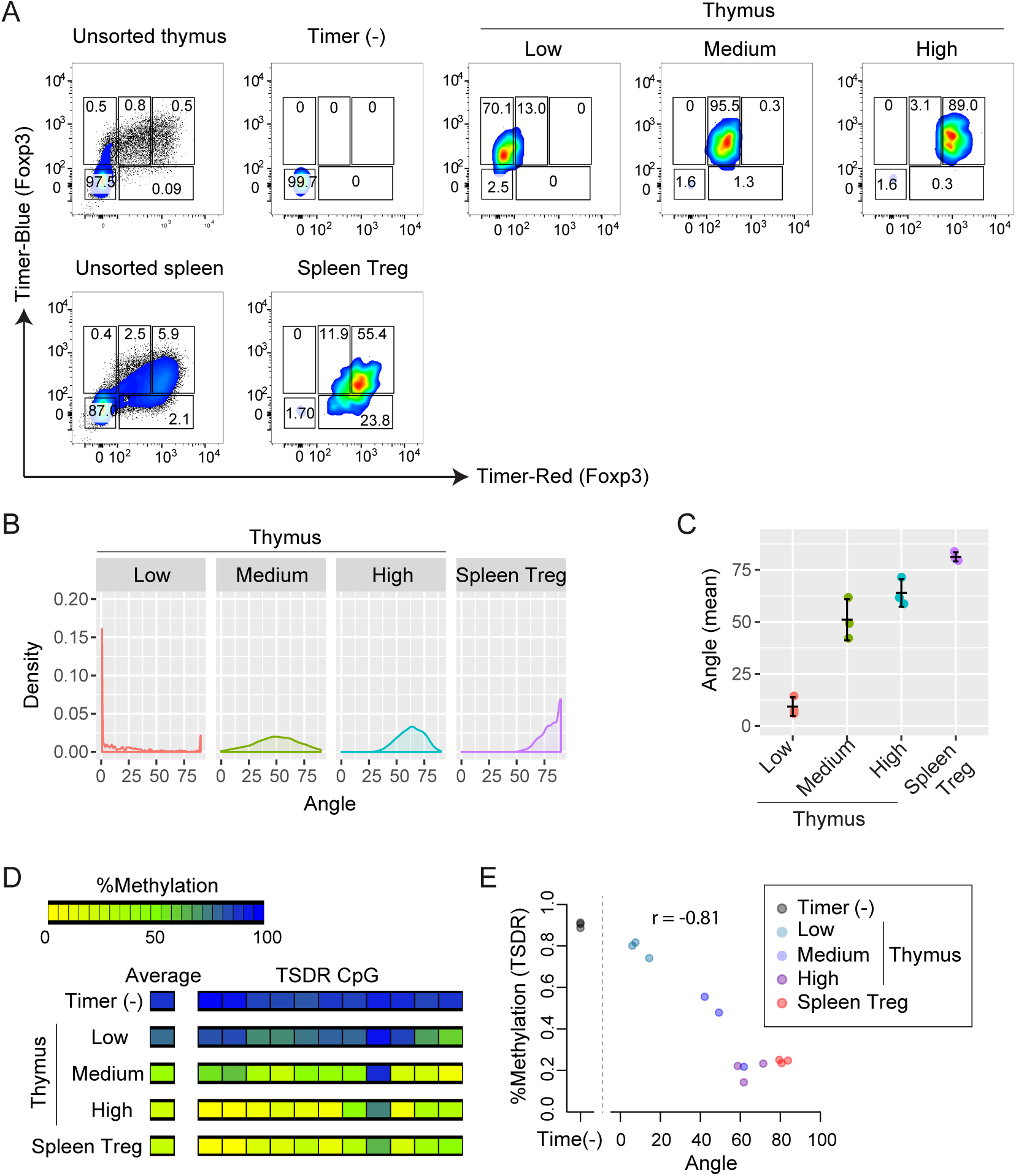
*Foxp3-Tocky* reveals *in vivo* dynamics of demethylation of the *Foxp3* gene. Thymic cells, or splenic Treg, were sorted according to increasing Tocky-time (i.e. Timer-Angle) in Blue+ cells. Displayed are **(A)** flow cytometry plots of Blue vs. Red fluorescence, **(B)** Timer Angle density plots, or **(C)** Mean Timer Angle in the sorted T-cell populations, n=3. DNA was extracted for methylation analysis, and thymic samples were compared to Timer (-) and splenic Treg samples **(D)** Heatmap showing average TSDR methylation rates of flow sorted T-cell subsets. **(E)** Average TSDR methylation rates plotted against mean Timer Angle in sorted T-cell subsets sorted, n = 3 mice for each subset. Spearman’s correlation coefficient is shown. Data were combined from two experiments.

Collectively, the *Foxp3*-Tocky reporter successfully identified newly generated Treg in vitro and in vivo, and also ordered cells from new to relatively aged ones (by the time domain analysis), demonstrating the general applicability of the Tocky system to studies of cellular biology and immunology.

## Discussion

Here we have established two uses of the Tocky system. First, the time domain analysis of Tocky allows investigators to determine the relative temporal order of the molecular events following the activation of key signalling pathways, and thereby provides a new way to identify immediate precursors and relatively mature cells. Previous studies using Fluorescent Timer proteins identified newly generated cells, as they expressed solely immature fluorescence without mature fluorescence (Miyatsuka et al., 2011; Miyatsuka et al., 2014; Terskikh et al., 2000). In this study, in addition to the rigorous determination of New cells (Blue^+^Red^-^), we established a quantitative method to measure the relative age of cells which are maturing in vivo. Thus, we determined the temporal sequences of molecular events during thymic Treg differentiation after receiving TCR signals using *Nr4a3*-Tocky, and identified the major precursor of these cells as the earliest cells among the presumptive precursor populations. In addition, the time domain analysis has allowed us to simultaneously investigate the temporal dynamics of the demethylation process and the transcriptional activities in vivo, identifying and characterising the earliest stages of Foxp3^+^ Treg differentiation. Instead of using a surface marker as an indicator of developmental stage (Toker et al., 2013), we directly investigated the time domain of *Foxp3* transcription and thereby revealed the in vivo dynamics of the demethylation of the *Foxp3* enhancer region: the demethylation process is initiated after *Foxp3* transcription started and it becomes the most active when the transcriptional activity is highly sustained. This means that the Tocky system provides an unprecedented means to investigate the in vivo regulation of molecular mechanisms.

Second, the frequency domain analysis of the Tocky system can identify cells that are undergoing repeated signaling processes. Using *Nr4a3-Tocky*, we showed that thymic T cells that have received sustained TCR signals differentiate into Treg. Consistent with these findings, thymic Foxp3^+^ cells show slower and more confined migration, compared to other thymic T cells by two-photon microscopy (Le Borgne et al., 2009). These differentiating Foxp3^+^ cells may integrate TCR and other signals from thymic epithelial cells and other APCs. In addition, using Nr4a3-Tocky, tissue-infiltrating antigen-specific T cells also accumulate in the Persistent locus, indicating that they frequently interact with cells presenting their cognate antigens. This is also compatible with the findings by intravital microscopic analysis of TCR transgenic T cells that showed that antigen-specific T cells decrease velocity when interacting with antigens (Bousso, 2008). Since the affinity of TCR-ligand interactions determines the dynamics of proximal TCR signalling molecules (Stepanek et al.), future studies should investigate whether and how different TCR affinities are translated into different dynamics of transcription of downstream transcription factors, including Nr4a3. In addition, further studies are required to investigate the temporal dynamics of antigen-specific T cell responses in different contexts such as other autoimmune diseases, allergy, infections, and vaccination. Nevertheless, *Nr4a3-Tocky* enables the investigation of the antigen-specific response of T cells within polyclonal repertoires, revealing in vivo T cell responses at the single cell level, which may be effective in evaluating the effects of immunotherapy on T cell responses.

In addition, using Nr4a3-Tocky, self-reactive T cells are identified as cells that receive infrequent TCR signals in the periphery. The spontaneous TCR signals in self-reactive T cells are historically defined as “tonic TCR signals” (Stefanova et al., 2002), although their temporal dynamics were unknown. Using Nr4a3-Tocky, most of self-reactive T cells are mainly at the Arrested locus, and therefore the interval of signals is considered to be several times longer than the half-life of Blue fluorescence (~4 h). Further study is required to elucidate the mechanism of the infrequent TCR signals in self-reactive T cells and whether and how TCR and other signals differentiate self-reactive T cells into pathogenic T cells in autoimmune conditions. It is of interest how different frequencies of TCR signals result in the activation of different transcriptional mechanisms.

In general, persistent signals may have distinct biological roles compared to transient signals (Yosef and Regev, 2011), and the Tocky system is an effective tool to investigate these dynamics. For example, sustained TLR4 signalling induces *Il6* transcription effectively, in contrast to transient activities (Litvak et al., 2009); sustained DNA damage, but not transient damage, activate p53 and induces p21 expression and cell cycle arrest (Loewer et al., 2010). To date, such studies used in vitro time course analysis and/or mathematical modelling to analyse the sustained dynamics of transcription. The Tocky system will benefit studies in cell signalling by providing a means to directly identify and isolate cells receiving persistent signals.

Thus, the Tocky system can be used to dissect the temporal dynamics of cellular differentiation and activation of individual cells by analysing their time- and frequency domains, providing a measurement of ‘Tocky-time’ (**Supplementary Fig. 4**), which is in a non-linear relationship with ‘real’ time, and represents a relative chronological readout for events occurring following a cellular differentiation cue. It is of note that the time and frequency domains of the Tocky-time are merged at the Persistent locus, and further mathematical approaches to understand the relationship between these two domains are anticipated. The advantages of the Tocky system over other gene reporters are summarised in **Supplementary Table 1**. In the future Tocky mice for key transcription factors and genes will be promising tools to reveal the in vivo dynamics of gene transcription and cellular differentiation (e.g. Bcl6, for T-follicular helper cells; Rag2, for TCR recombination; and, Oct4, for stem cell-ness), which cannot be investigated otherwise.

In summary, we have established the Tocky system as a pioneering tool to investigate cellular activation and differentiation and gene dynamics in vivo, which will facilitate studies in cell biology disciplines including immunology, developmental biology and stem cell biology.

## Materials and Methods

### Transgenesis and mice

The BAC clones RP23-122N18 and RP23-330D17 were obtained from the BACPAC Resources Center at Children’s Hospital Oakland Research Institute (CHORI), and were used for generating *Nr4a3*-Tocky and *Foxp3*-Tocky, respectively. BAC DNA was modified by the BAC recombineering approach using the *SW106* strain bacteria (Warming et al., 2005). Two independent lines were established for both of the *Nr4a3*-Tocky and *Foxp3*-Tocky transgenic reporter strains. Both lines exhibited highly similar phenotypes and frequencies of Timer^+^ cells.

We used a Timer knock-in knock-out approach for BAC transgenic reporter constructs. Precisely, for *Nr4a3*-Tocky, the first coding exon of the *Nr4a3* gene in RP23-122N18 was targeted and replaced with the transgene cassette containing the *Timer (Fast-FT)* gene (Subach et al., 2009), a poly-A tail and a floxed neomycin resistance gene *(neo)*. For *Foxp3*-Tocky, the first coding exon of the *Foxp3* gene in RP23-330D17 was targeted and replaced by the same transgene cassette. Subsequently, *neo* was excluded by arabinose-inducible Cre expression in SW106 (Warming et al., 2005). BAC DNA was purified by the NucleoBond Xtra Midi kit (Macherey-Nagel), and microinjected into the pronucleus of one-cell embryos from C57BL/6 mice under the approval by the Gene Recombination Experiments Safety Committee of Kyoto University. Founders were screened by genomic PCR, and transgene-positive founders were mated with wild-type C57BL/6 mice. F1 mice were screened by flow cytometry for Timer expression, and subsequently bred to homozygosity with *Foxp3^IRES-GFP^* mice (B6.Cg-*Foxp3^tm1Mal^/J*, Jackson Laboratories, #018628), to generate *Nr4a3*-Tocky:*Foxp3^IRES-GFP^* double transgenic mice. *OT-II Nr4a3*-Tocky: *Foxp3^IRES-GFP^* mice were similarly generated by crossing *Nr4a3*-Tocky, B6.Cg-*Tg(TcraTcrb)425Cbn/J*, and B6.Cg-*Foxp3^tm1Mal^/J*. MHC Class II ^ko^/^ko^ (B6.129S2-H2dlAb1-Ea/J, Jackson Laboraotries #003584) and congenic CD45.1 (B6.SJL-Ptprca Pepcb/BoyJ, Jackson Laboratories #002014) were also used. All animal experiments were performed in accordance with local Animal Welfare and Ethical Review Body at Imperial College London (Imperial) and University College London (UCL), and all gene recombination experiments were performed under the risk assessment that was approved by the review board at Imperial and UCL.

### *In vitro* Treg polarisation and mature Treg culture

CD4+CD44^lo^Foxp3^-^ naïve T-cells from *Foxp3*-Tocky mice were isolated by cell-sorting and 1 × 10^5^ cells cultured on anti-CD3 (clone 1452C11, 2 μg/ml) and anti-CD28 (clone 37.51,10 μg/ml; both eBioscience)-coated 96 well plates (Corning) in the presence of 100 U/ml rhIL-2 (Roche) and 2 ng/ml rhTGFβ (R&D) for 0 - 48 h in a final volume of 200 μL RPMI1640 (Sigma) containing 10% FCS and penicillin/streptomycin (Life Technologies).

Mature Foxp3^+^ Treg from *Foxp3*-Tocky mice were isolated by cell-sorting and 1 × 10^5^ cells cultured on anti-CD3 (clone 145.2C11, 2 μg/ml) and anti-CD28 (clone 37.51,10μg/ml; both eBioscience)-coated 96 well plates (Corning) in the presence of 100 U/ml rhIL-2 (Roche) for 0 - 48 h in a final volume of 200 μL RPMI1640 (Sigma) containing 10% FCS and penicillin/streptomycin (Life Technologies).

### Activation of T and B cells *in vitro* and the analysis of the half-life of Blue-fluorescence

Splenocytes (4 × 10^5^/ well) were cultured on 96 well U bottom plates, coated with 2 μg/ml anti-CD3 (145.2C11, eBioscience) and/or 10 μg/ml antiCD28 (37.51, eBioscience) for 20 h. Cells were then harvested and replated in the presence of 100 μg/ml cycloheximide (Sigma). At various time points, cells were stained with CD4 and CD8 antibodies and Timer-Blue and -Red fluorescence measured by flow cytometry. In some cultures, 100 U/ml rhIL-2 (Roche) was added. For polyclonal activation of B cells, 10 μg/ml of F(ab')2-goat anti mouse IgM (Life Technologies) was added to cultures of splenocytes for 20 h, Cells were then harvested and re-plated in the presence of 100 μg/ml cycloheximide (Sigma). At various time points, cells were stained for CD19 and CD3, and CD3-CD19+ B cells analysed for Timer-Blue and -Red fluorescence.

### *In vitro* T cell activation of OT-II *Nr4a3-Tocky* T cells

CD4^+^ T cells from OT-II *Nr4a3*-Tocky:*Foxp3^GFPKI^* mice were isolated by immunomagnetic cell separation, (StemCell Technologies) and 2 × 10^5^ cells cultured with 3 × 10^5^ (2:3) CD90.2-depleted splenocytes in the presence of 1, 10, 100 or 1000 nM Ova_(323–339)_ peptide (Sigma) on 96 well plates (Corning) in a final volume of 200 μL RPMI1640 (Sigma) containing 10% FCS and penicillin/streptomycin (Life Technologies) and 55 μM beta mercaptoethanol (Gibco) for the stated time periods. At 24 h, some cells were washed three times and recultured on a fresh plate in the presence of 40 μg/ml anti-MHC Class II (clone M5/114, BioXCell) for a further 8–24 h before analysis. The stimulation was terminated in the same manner for periodic stimulation of OT-II Nr4a3.

### Confocal Microscopy

Confocal microscropy analysis of stimulated OT-II: *Nr4a3*-Tocky T cells was performed using Cytospin as previously described (Ono et al., 2007). Briefly, Cytospin slides were prepared with stimulated T cells and fixed by 2% paraformaldehyde for 10 min at room temperature. After washing with PBS, fluorescence images were obtained by the confocal system, the Zeiss LSM510 invert (Carl Zeiss, Jena, Germany) using a Plan-Apochromat 20x/0.8 objective. The LSM image files were converted into the TIFF format and further analysed by Fiji (Schindelin et al., 2012). For the quantitative analysis of fluorescence intensities in single cells, individual cells were manually identified as Region of Interest (ROI) using the light field Differential interference contrast (DIC) images. It is of note that GFP+ Foxp3^+^ cells were very rare in the stimulated OT-II *Nr4a3*-Tocky cells, and therefore we used GFP images as a dump excluding cells with high autofluorescence (as shown in **Supplementary Fig. 2** by arrowheads). The measurement of mean fluorescence intensities were obtained by redirecting it to thresholded greyscale images for Timer-Blue and Timer-Red fluorescence. The same threshold value was applied to all the images from the same channel. Mean fluorescence intensities were logged to produce histograms and 2-dimensional plots. Colour images were enhanced for brightness by applying the same linear adjustment to all the images from the same channel.

### Experimental Autoimmune Encephalomyelitis

EAE was induced by subcutaneous injection of 200 µg Myelin Oligodendrocyte Glycoprotein (MOG_35–55_, Sigma) emulsified in Freund’s adjuvant (Sigma) containing 4 mg/ml heat killed mycobacteria (Invivogen). Control mice received CFA alone, emulsified with sterile water. On Day 0 and Day 2, mice received 200 ng i.p. of pertussis toxin (Calbiochem). Mice were monitored for the development of clinical symptoms: 0 = no clinical symptoms;0.5: Tip of tail is limp; 1: limp tail; 1.5: limp Tail and hind leg inhibition; 2: limp tail and weakness of hind legs: 2.5: limp tail and dragging of hind legs; 3: limp tail and paralysis of hind legs. The severity limit of the protocol was 3.

For isolating cells from the spinal cord, mice were culled and the left ventricle perfused with ice cold PBS. Spinal cords were removed and forced through a 70 μm cell strainer. CNS lymphocytes were separated from myelin using a (30/70%) Percoll gradient (Sigma). Mononuclear cells were removed from the interphase, washed and resuspended in 10% RPMI for labelling.

### Tetramer staining

dLN or CNS mononuclear cells isolated by Percoll gradient were incubated at 37°C for 15 minutes in the presence of 50 nM Dasatinib (Lissina et al., 2009). 1 in 100 dilution of APC-labelled mouse I-A^b^ MOG peptide 38–49 (GWYRSPFSRVVH, I-Ab MOG_38–49_, NIH Tetramer Core) was added to the cells and further incubated for 30 minutes at 37°C. Cells were then washed and stained for viability dye and surface markers on ice.

### Flow cytometric analysis and cell sorting

Following spleen or thymus removal, organs were forced through a 70 µm cell strainer to generate a single cell suspension. For splenocyte preparations a RBC-lysis stage was employed. Staining was performed on a V-bottom 96-well plate, or in 15 ml falcon tubes for cell sorting. Analysis was performed on a BD Fortessa III instrument. The blue form of the Timer protein was detected in the blue (450/40 nm) channel excited off the 405 nm laser. The red form of Timer protein was detected in the mCherry (610/20) channel excited off the 561nm laser. For all experiments a fixable eFluor 780-fluorescent viability dye was used (eBioscience). The following directly conjugated antibodies were used in these experiments: CD3 PerCPcy5.5 (clone 145.2C11, TONBO) CD4 APC (clone RM4-5, eBioscience), CD4 Alexa-fluor 700, CD4 BUV395 (clone GK1.5 BD Biosciences), (clone RM4-5, Biolegend), CD8 PE-Cy7 or APC (clone 53-6.7, Biolegend), CD8 BUV737 (clone 53-6.7, BD Biosciences) TCRβ FITC & Alexafluor 700 (clone H57-597, Biolegend), TCRβ BUV737 (clone H57-597, BD Biosciences), CD19 APC (clone 6D5, Biolegend), CD25 PerCPcy5.5 (PC61.5, eBioscience) or PE-Cy7 (PC61.5, Tombo Bioscience), CD44 APC (clone IM7, eBioscience) or Alexafluor 700 (clone IM7, Biolegend), CD69 APC (H1.2F3, eBioscience) and Foxp3 APC (clone FJK-16s, eBioscience), CD45.1 PE-Cy7 (clone A20, Biolegend), CD45.2 APC (clone 104, Biolegend), CD45RB PE-Cy7 (clone C363-1**6A)**. For *Foxp3-Tocky* validation, RBC-lysed splenocytes were stained with eFluor 780 viability dye, before staining for APC-conjugated CD4 and FITC-conjugated TCRβ. Live CD4^+^TCRβ^+^ were gated and then sorted into four fractions: Timer(Blue^+^Red^−^), Timer (Blue^+^Red^+^), Timer(Blue^−^Red^+^), Timer(Blue^−^Red^−^). Cells were then fixed and labelled with APC-conjugated Foxp3 using the eBioscience Foxp3 fixation and permeabilisation kit according to the manufacturer’s instructions. In some experiments, cells were labelled with a eFluor 670 proliferation dye (eBioscience), at a concentration of 5 μM for five minutes at room temperature.

### Adoptive Transfer Experiments

CD4^+^CD25^-^CD44^lo^Foxp3^-^T-cells from OT-II *Nr4a3*-Tocky mice were isolated by cell sorting on FACS Aria III, and between 1-5 M cells were adoptively transferred i.p. to congenic CD45.1 recipients, which were fed ovalbumin (Sigma) at a concentration of 1.5% in water. For MHC Class II Knock out experiments, CD4+ T-cells were isolated by immunomagnetic selection (Stem Cell Technologies) and 3–5 M cells injected i.p. into congenic (CD45.1) MHC Class II ^ko^/^ko^ mice. MHC Class II ^ko^/^wt^ mice were used as controls. Adoptively transferred T-cells were defined as CD45.1^-^CD45.2^+^CD4^+^TCRβ^+^ cells.

### DNA methylation analysis

DNA was extracted from sorted samples using Qiagen DNeasy kit according to the manufacturer’s instructions. 100 ng of DNA was bisulfite treated using the Epitect Bilsulfite Kit (Qiagen), and used as a template for amplification of the TSDR. TSDR was amplified using the following primers (Foxp3 TSDR For: TTTGAATTGGATATGGTTTGT; Foxp3 TSDR Rev: AACCTTAAACCCCTCTAACATC (Floess et al., 2007)) and cycling conditions: 94 °C 1min followed by 40 cycles of 94 °C 15 s / 54 °C 30 s / 68 °C 30 s and a final extension phase of 68 °C for 15minutes. PCR amplicons were purified and then underwent Sanger Sequencing using the reverse primer by Source BioScience. Sequence traces and CpG methylation rates were analysed and determined by ESME software (Epigenomics) (Lewin et al., 2004).

### Canonical Correspondence Analysis

The gene signature and sample scores in Supplementary Fig. 1 were calculated by a cross-dataset analysis using CCA (Ono et al., 2014). Briefly, the expression data of GSE15907 (Painter et al., 2011) was regressed onto the log2 fold change of activated CD4^+^ T cells (2 h after activation) and naïve T cells from GSE48210 (Li et al., 2013) as the explanatory variable, and Correspondence Analysis was performed for the regressed data and the correlation analysis was done between the new axis and the explanatory variable. CCA was performed by the CRAN package *vegan* as previously described (Ono et al., 2014). The analysis was undertaken using only transcription factor genes, which were selected by the Gene Ontology database by including the genes that are tagged with GO:0003677 (DNA binding) and GO:0005634 (nucleus) and not with GO:0016020 (membrane) using the Bioconductor package *GO.db*.

### Timer data analysis

#### 1. Overview of Timer data analysis

Timer data analysis is composed of the following three steps: (1) Data preprocessing and scaling / normalisation of Blue and Red fluorescence data; (2) Trigonometric data transformation of Blue and Red fluorescence, which transform data into Timer-Angle and -Intensity using the polar coordinate; and (3) Data export, statistical analysis and visualisation. Timer data analysis will be performed by importing those csv files into *R* (R Core Team, 2016). A series of data preprocessing, normalisation and transformation results in Timer normalised data, which are further analysed for statistics, visualisation, and quality control (QC) processes. Finally the code exports these QC and statistical results, the cell number data of total cells and Timer^+^ cells in each file, and a matrix containing Timer-Angle and -Intensity data and all the fluorescence data for individual cells in each file (**Supplementary Fig. 5A)**.

#### 2. Data import

Flow cytometric data will be gated for T cell populations (e.g. CD4+ T cells) by an external programme such as FlowJo or the Bioconductor package *flowCore*, and be batch exported as csv files including negative control file and FSC control file (see below). The R codes will import all the files and produce and work on the dataset as an R object.

#### 3. Data preprocessing

Data preprocessing for Timer data analysis is composed of the following three steps: (1) FSC correction; (2) data thresholding; and (3) data normalisation.

##### 3.1. FSC correction

Our investigations showed that Blue autofluorescence increases as ForwardScatter (FSC) increases, which indicates that larger cells have higher Blue signals (**Supplmentary Fig. 5B)**. This effect was more remarkable in Blue than Red, which can result in underestimation of Timer-Angle, for example, in activated T cells, which have larger cell sizes. Accordingly, the code incorporates the function to perform FSC correction by applying a linear regression to FSC (*x*) and either Blue (*b*) or Red (*r*):

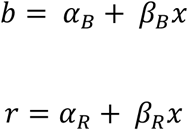

Thus, FSC-corrected Blue *(b****_c_***) and FSC-corrected Red *(r****_c_***) are obtained by:

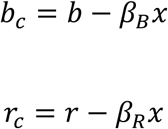

##### 3.2. Data thresholding

The instrument setting of flow cytometry is determined by manually adjusting the amplification parameters, so that the autofluorescence signals of negative cells have positive values, and that the threshold value for positive signals can be visually determined. In the standard practice of flow cytometric analysis, the threshold value for each fluorescence is determined using a negative control cell sample, so that a certain small proportion of cells are identified as positive, for example 0.5% of the parent population (Fujii et al., 2016). These negative signals are problematic for trigonometric data transformation, because Blue^low^Red^-^ and Blue^-^Red^low^ cells can have various Timer-Angle values, which is biologically meaningless (**Supplementary Fig. 5C)**. Accordingly, the negative signals of FSC-corrected Blue (*b****_c_***) and FSC-corrected Red data *(r****_c_***) will be collapsed using the threshold value *n****_B_*** and *n****_R_***, respectively.

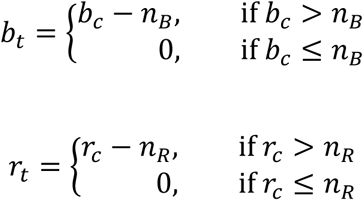

In **Supplementary Fig. 5C**, Time-Angle calculation with or without Data thresholding are compared on uniformly distributed cell data. Here it is assumed that each cell has a discrete integer value between 0 and 100 for Blue and another value for Red (i.e. there are 10201 cells in the Blue-Red plane), and that the barycentre of negative control cells is (10, 10). When data were processed by Blue-Red normalisation without applying data thresholding, cells can have various Timer-Angle values around the barycentre of negative control cells, and thus, cells in the negative quadrant gate (the lower left quadrant of pink lines) can have all the different Timer-Angle values (**Supplementary Fig. 5C**, top left) and have all the Timer loci (**Supplementary Fig. 5C**, top right). These variations in Blue^-^ Red^-^ cells are biologically meaningless. In **Supplementary Fig. 5C, bottom left**, data were firstly thresholded (at Blue = 18, Red = 18 in this example), and thereafter Blue and Red data were normalised. The thresholding removes all the Blue^-^Red^-^ cells. All the Blue^-^ Red+ cells have Timer-Angle value 90, while all the Blue^+^ Red^-^ have Timer-Angle value 0.

##### 3.3. Log transformation

After thresholding negative signals, Timer fluorescence data are log transformed by the function *log*(*x +* 1).

##### 3.4. Blue-Red normalisation

The autofluorescence data of Blue and Red fluorescence in negative control sample will be used to estimate the deviation of Blue and Red signals in each dataset. Assuming that the standard deviation of Blue and Red fluorescence in negative control is *σ****_Β_*** and *σ****_R_***, normalised Blue and Red data *b****_n_*** and *r****_n_*** are obtained as follows:

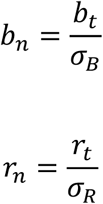

#### 4. Trigonometric data transformation

Lastly, using a trigonometric function, normalised Blue and Red data will be transformed into Timer-Angle and Timer-Intensity data. Timer-Angle *θ* is defined as the angle from the normalised Blue axis and has the value between 0 and 90 degree, or [0, 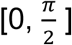]. Timer-Intensity *δ* is the distance from the origin of normalised Blue and normalised Red axes (**Supplementary Fig. 5D)**. By definition, cells in the origin (0, 0) are negative for Timer protein, and are removed from further analysis. Thus, the R code automatically identifies Timer^+^ cells.

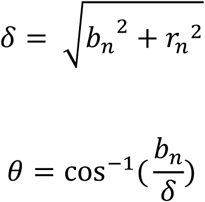

#### 5. Timer Locus analysis

Timer Locus analysis allows the translation of Timer data into transcriptional dynamics and the biological interpretation of individual cells (**Supplementary Fig. 5E)**. The 5 Timer loci are designated as follows: New = 0°; New-Persistent transition (NP-t) = (0°, 30°); Persistent [30°, 60°]; Persistent-Arrested transition (PA-t) [60°, 90°]; and Arrested = *90°* (**Supplemntary Fig. 5F)**. Nonparametric tests for Timer locus were performed by the *CRAN* package. Kruskal test was applied to data with more than 2 experimental groups, and subsequently post-hoc Dunn’s test was applied.

## Computer simulation of Timer fluorescence

Computer simulation of GFP and Timer expression was performed by time course analysis of a series of first-order equations involving the function for transcript (mRNA) *X* and translated Timer proteins including the colourless-form *C*, the blue-form *B*, the intermediate-form *I* (which does not emit fluorescence), and the red-form *R*, or translated GFP protein *G*, and the decayed protein *D*. Thus, a liner kinetic model for Timer protein was constructed as previously reported (Subach et al., 2009):

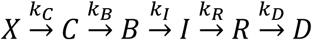

And for GFP:

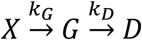

The amounts of blue- and red-form proteins are

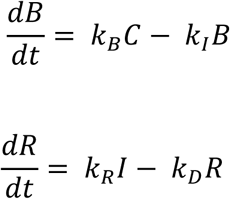

Similarly, the amounts of other intermediate form proteins were modelled in a similar manner, and time course analysis was performed by the CRAN package *deSolve*. Analytically, the amounts of blue- and red-form proteins are as follows:

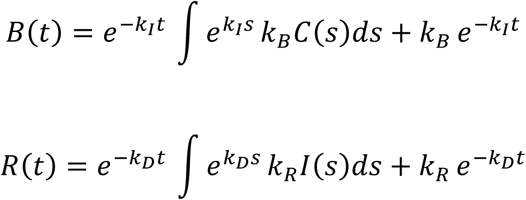

Thus, blue- and red- form proteins are dependent on the supply from their precursor proteins, and when transcription is terminated and no precursor protein is supplied, they are expected to exponentially decay at the rate *k_B_* and *k_R_*. For **Fig. 1C**, the parameters from the preceding study (Subach et al., 2009) were used to perform time course analysis. For GFP, the translation rate was set to be the same with Timer, *k_G_ =* 3.0 × 10^−1^*/h*, while the decay rate to be *k_E_ =* 1.3 × 10^−2^*/h*, as previously reported (Sacchetti et al., 2001). Density plots for the distribution of Timer-Angle (**Fig. 1G)** were generated by testing different frequencies and intermittent expressions, assuming that signals are conveyed to cells for 7 days by the temporal dynamics shown in each figure, and that each cells is at a random time point in this duration.

## Statistical analysis and data visualisation

Statistical analysis was performed on R or Prism 6 (Graphpad) software. Percentage data for Timer^+^ and Timer locus analysis was analysed by Mann-Whitney U test or Kruskal-Wallis test with Dunn’s multiple comparisons using the CRAN package *PMCMR*. Samples with fewer than 18 Timer^+^ cells were not included in the analysis. Student’s t-test was used for comparison of two means. For comparison of more than two means, a one-way ANOVA with Tukey’s post-hoc test was applied using the CRAN package *Stats*. Scatterplots were produced by the CRAN packages *ggplot2* and *graphics*. Dose response data in **Fig. 3G** were fitted to a dose-response curve using the three-parameter log-logistic function of the CRAN package *drc* as previously described (Yoshioka et al., 2012). All computations were performed on Mac (version 10.11.6). Adobe Illustrator (CS5) was used for compiling figures and designing schematic figures. Variance is reported as SD or SEM unless otherwise stated. * p<0.05, ** p<0.01, *** p<0.001, **** p<0.0001.

## Data and code availability

All R codes are available upon request. Data will be made available upon reasonable requests to the corresponding author.

## Supplementary Materials

Supplementary Fig. 1: CCA identifies *Nr4a3* as a downstream target of TCR signalling.

Supplementary Fig. 2: Confocal microscopy analysis of OT-II Nr4a3-Tocky T cells stimulated at various frequencies.

Supplementary Fig. 3: Steady state TCR signalling is restricted to Memory-like T-cells in vivo and dependent on MHC Class II.

Supplementary Fig. 4. Summary of Tocky technology.

Supplementary Fig. 5. Timer data analysis methods.

Supplementary Table 1. Differences between Tocky, Fate-mapper, and GFP/FP reporters.

## Author Contributions

M.O. conceived the Tocky technology. M.O. designed and generated transgenic constructs. S.K., H.M., and M.O. established the transgenic founders. S.K., H.M., P.P.M, and D.B screened mouse lines. D.B., T.C, and M.O conceived and designed immunological experiments. D.B., P.P.M, A.P., and C.D. performed animal experiments. D.B., P.P.M., A.P., E.M., and M.L. performed in vitro experiments. M.O. conceived Timer data analysis, wrote computational codes and performed bioinformatics analysis and data visualisation. D.B., T.C., and M.O. wrote the manuscript.

## Acknowledgements

We thank for their kind support at the Flow Cytometry facility, Dr. Ayad Eddaoudi and Ms Stephanie Canning (University College London), and also Ms Jane Srivastava, Ms Catherine Simpson, and Ms Jess Rowley (Imperial College London). We thank Dr Andreas Bruckbauer for his technical support at the Facility for Imaging by Light Microscopy, Imperial College London. We thank Prof. Anne Cooke (University of Cambridge), Prof. Charles R. M. Bangham (Imperial College London), Prof. Fiona Rawle (University of Toronto), Prof. Taku Okazaki (Tokushima University), Prof. Gloria Rudenko (Imperial College London) and Dr. Alastair Copland (St. George’s University London) for their feedback on the manuscript.

M.O. is a David Phillips Fellow (BB/J013951/2) from the Biotechnology and Biological Sciences Research Council (BBSRC). T.C. is supported by the Medical Research Council and Great Ormond Street Hospital Children’s Charity.

The authors declare that no conflicts of interest exist in relation to this manuscript.

---

**Supplementary Fig. 1:**
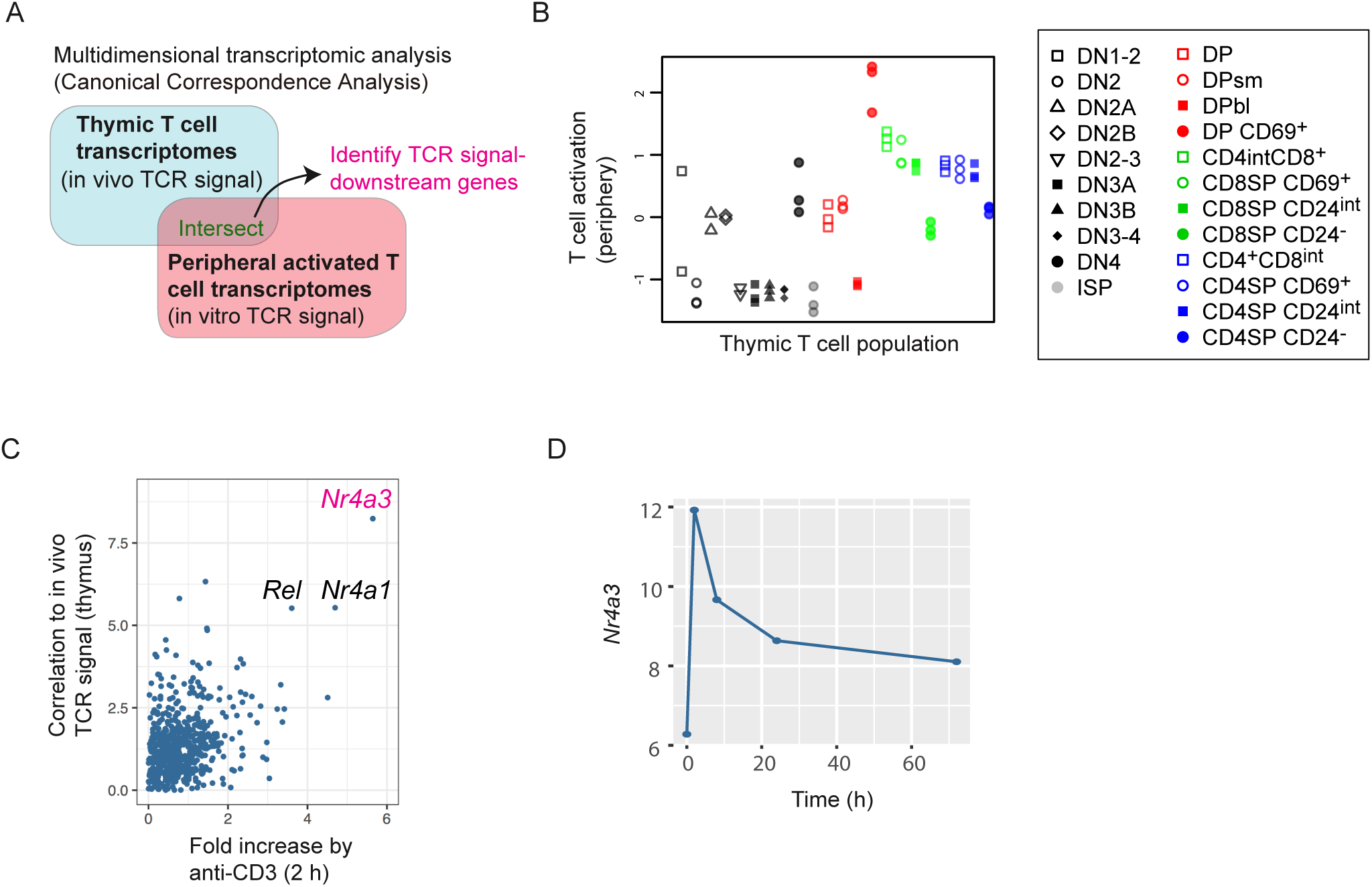
CCA identifies *Nr4a3* as a downstream target of TCR signalling. **(A)** Schematic showing rationale for identifying genes that are downstream of TCR signalling. Cross-dataset analysis of the transcriptomes of thymic T cell populations (in vivo signal, GSE15907) and those of TCR-signalled peripheral CD4+ T cells data (in vitro signal, GSE48210). CCA was performed to identify the candidate genes used by both thymic and peripheral T cells. **(B)** Thymic T cell population data was analysed by CCA using activated and resting T cells from GSE48210 as the explanatory variable. The output of CCA is composed of (1) cell sample score, (2) gene score, and (3) biplot value for explanatory variable, and thus allowing the cross-level analysis of cells, genes, and biological process. Thus, using the axis that is correlated with T cell activation, T cell activation scores of thymic T cell populations were specified and visualised. **(C)** Scatter plot showing the fold change increase by anti-CD3 stimulation and the correlation to the thymic T cell populations that received TCR signals. Cross-dataset analysis was performed by CCA of the transcriptome data from thymic T cell populations *(in vivo* signal, GSE15907) and peripheral CD4+ T cells that received *in vitro* anti-CD3 stimulation *(in vitro* signal, GSE48210), in order to identify candidate genes upregulated by TCR signals in both thymic and peripheral T cells. **(D)** Time course analysis of *Nr4a3* transcripts upon anti-CD3 stimulation (from GSE48210).

**Supplementary Fig. 2:**
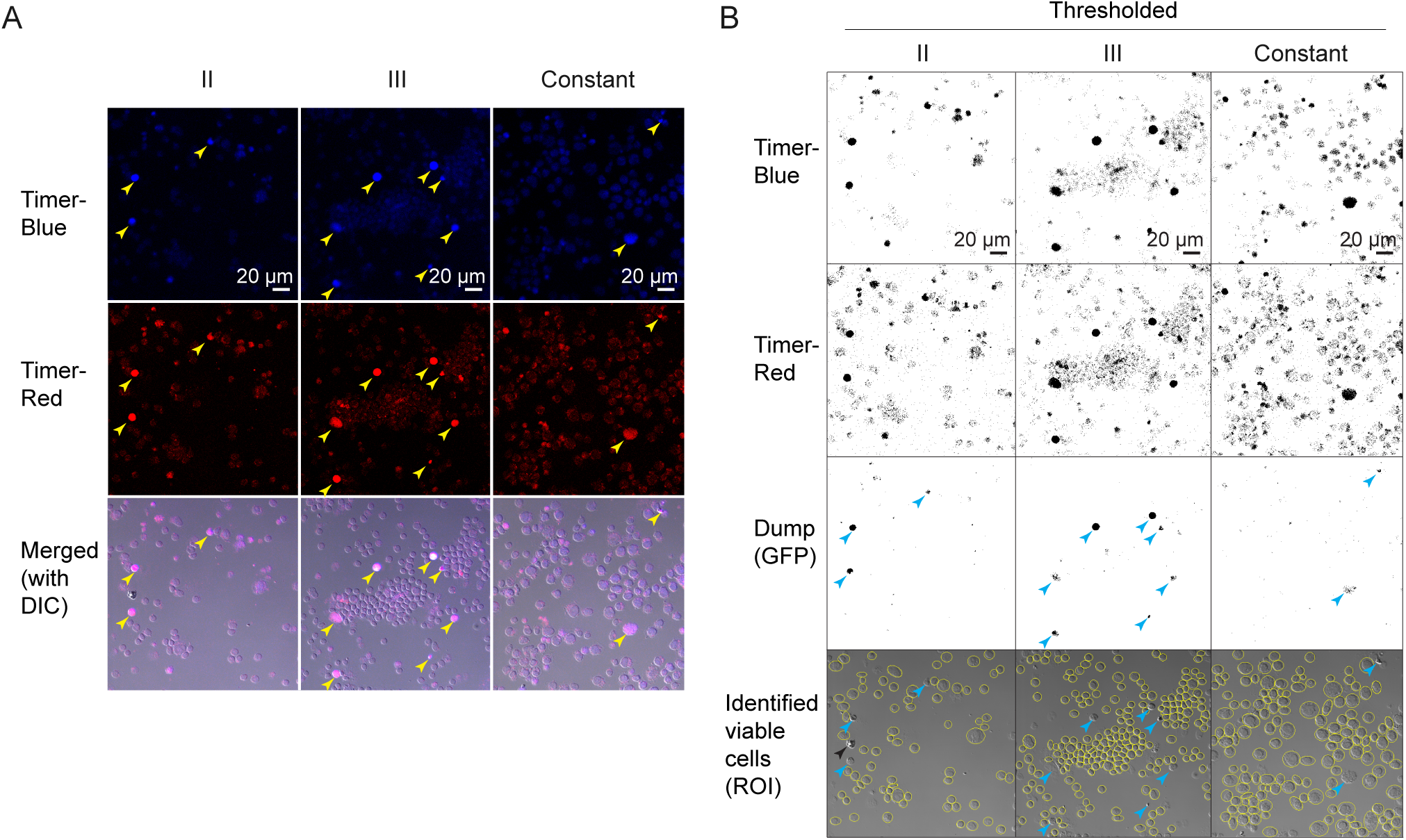
Confocal microscopy analysis of OT-II Nr4a3-Tocky T cells stimulated at various frequencies. T cells from OT-II Nr4a3-Tocky mice were stimulated with various frequencies. The Group II and III received 2 and 3 times 4-h stimulation by ova peptide within 48 h, respectively, and the Constant group received ova peptide stimulation continuously throughout the culture (see **Fig. 4**). Cells were attached to slides by Cytospin, fixed by paraformaldehyde and immediately analysed by confocal microscopy. **(A)** Confocal microscopy image of Timer-Blue, Timer-Red, and merged image (i.e. Timer-Blue and - Red with Differential interference contrast [DIC]). Arrowheads indicate apoptotic cells. **(B)** The images in A were thresholded using the same value for all the samples from the same channel. The GFP channel was used as a dump data for identifying apoptotic cells as those with high autofluorescence. DIC images were used to identify cells as Region of Interest (ROI), and the measurements in individual cells were redirected to thresholded greyscale images. Arrowheads indicate apoptotic cells identified by the dump GFP channel (coloured) or the morphology (black).

**Supplementary Fig. 3:**
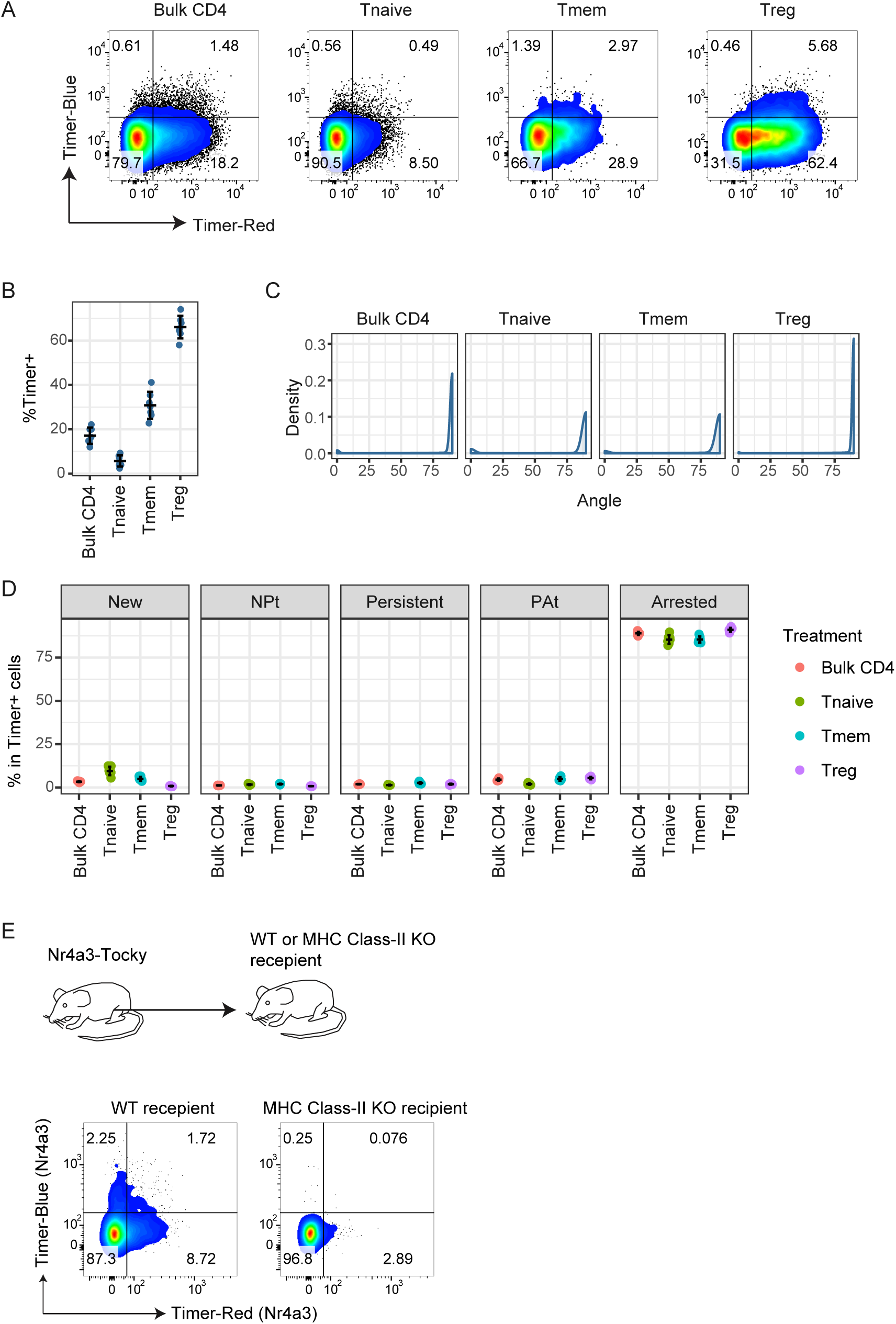
Steady state TCR signalling is restricted to Memory-like T-cells in vivo and dependent on MHC Class II. Nr4a3-Tocky mice (polyclonal repertoire) were analysed for Timer-Blue vs. Timer-Red fluorescence in CD4+ T-cells. **(A)** Flow cytometry plots showing Timer Blue vs. Timer Red fluorescence in the indicated populations from mLN. Bulk T-cells showed majority of Timer^+^ cells as Red+Blue-, meaning TCR signalling is relatively infrequent. CD4+ T-cells were sub divided into Naïve (CD4+CD44loCD25-Foxp3-) Memory (CD4+CD44hiCD25-Foxp3-) or Treg (CD4+Foxp3^+^). **(B)** Percentage Timer^+^ in the four T-cell subsets. **(C)** Density plot analysis reveals similar distribution of Timer Angles irrespective of CD4+ T-cell subset. **(D)** Timer locus analysis reveal Treg are greatly enriched with recent TCR-signalled cells. N=7, bars represent mean +/- SD. **(E)** CD4+ T-cells from Nr4a3-Tocky mice were adoptively transferred in to congenic WT or MHC Class II KO (I-Ab KO) mice. 9 days later transferred T-cells in the dLN were analysed for Timer-Blue vs Timer-Red expression. 2–5% of T-cells were receiving tonic signals in vivo, the majority of which were pure blue or pure red, indicating pulsatile signalling dynamics. Absence of Timer^+^ cells in MHC Class II KO mice demonstrates that Nr4a3-Tocky signalling is dependent on pMHC:TCR interactions in vivo.

**Supplementary Fig. 4.**
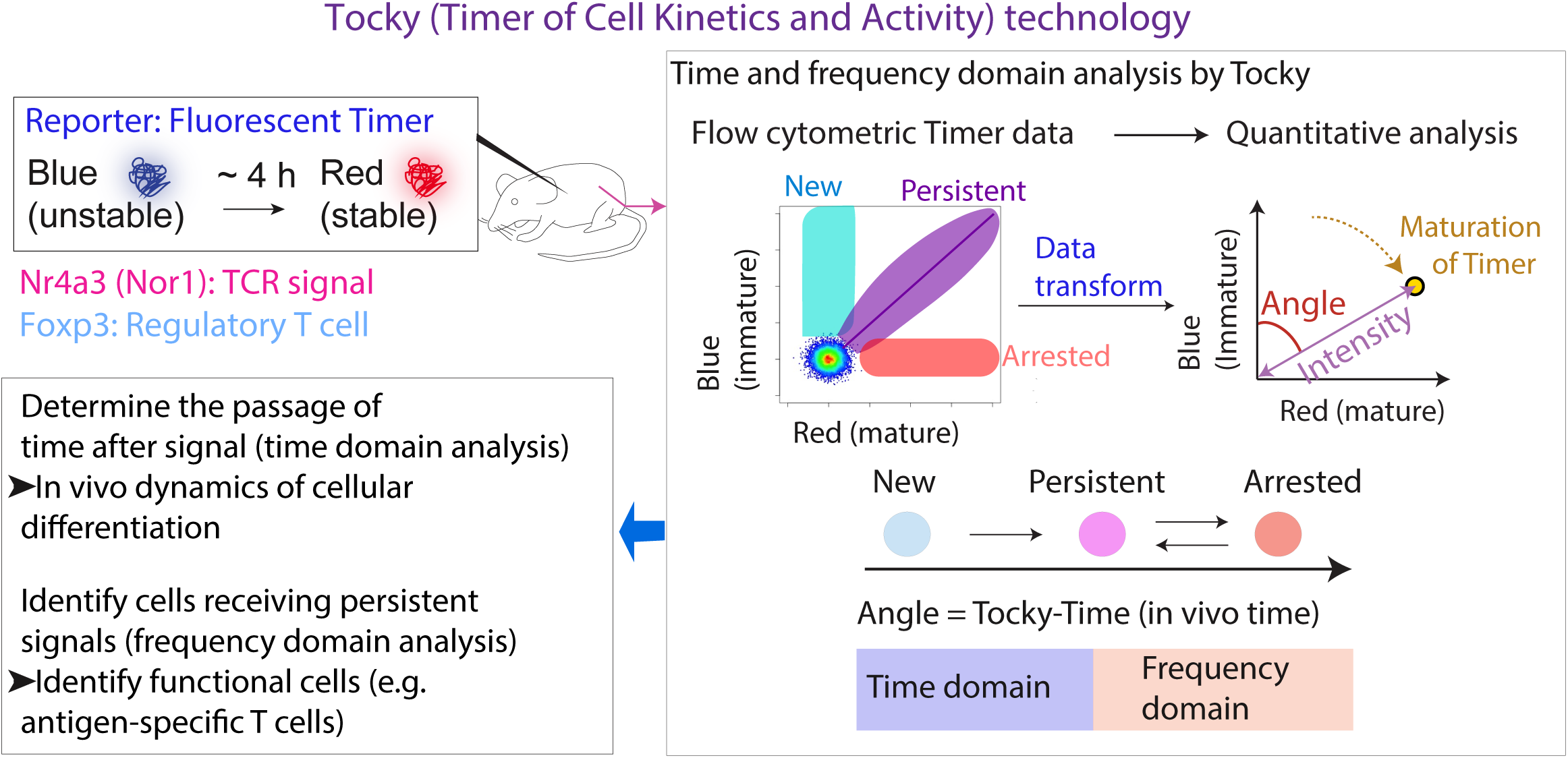
Summary of Tocky technology. Tocky system harnesses the power of Fluorescent Timer protein in combination with a computational algorithm to define the Time and frequency domains of cellular differentiation (Tocky-Time) using flow cytometry. This permits the determination of the relative passage of time following signalling cues, as well as identifying cells that are receiving continuous or very frequent differentiation signals.

**Supplementary Fig. 5.**
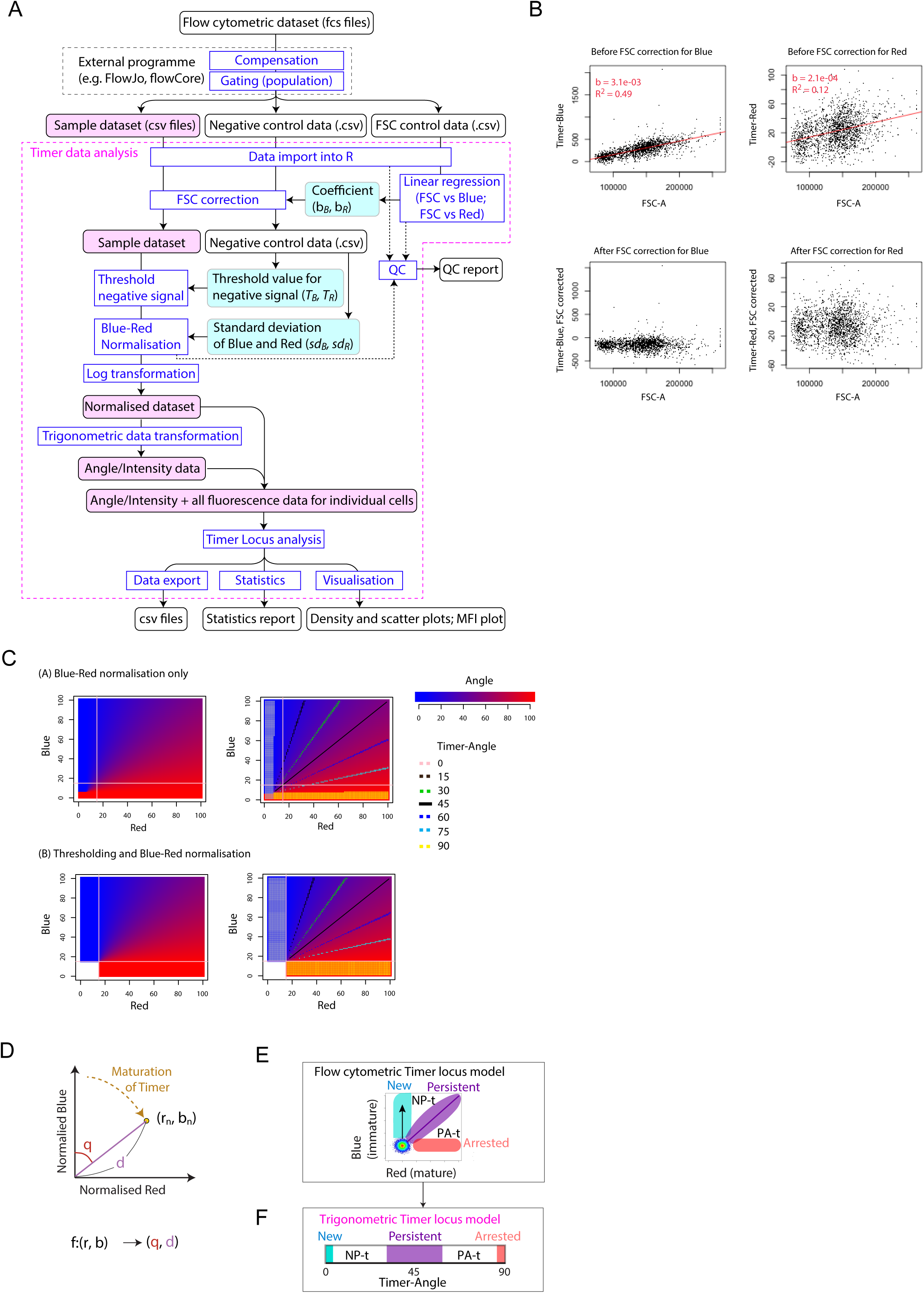
Timer data analysis. **(A)** Flow diagram detailing major components of Timer data analysis. **(B)** Flow cytometric plots showing linear regression of Timer Blue or Timer Red vs Forward Scatter (FSC). Bottom plots show Timer Blue or Timer Red following correction for FSC. **(C)** Timer-Angle calculation with or without data thresholding are compared on uniformly distributed cell data. Here it is assumed that each cell has a discrete integer value between 0 and 100 for Blue and another value for Red (i.e. there are 10201 cells in the Blue-Red plane), and that the barycentre of negative control cells is (10, 10). When data were processed by Blue-Red normalisation without applying data thresholding, cells can have various Timer-Angle values around the barycentre of negative control cells, and thus, cells in the negative quadrant gate (the lower left quadrant of pink lines) can have all the different Timer-Angle values (top left) and have all the Timer loci (top right). These variations in Blue^-^ Red^-^ cells are biologically meaningless. Data were firstly thresholded (at Blue = 18, Red = 18 in this example, bottom left), and thereafter Blue and Red data were normalised. The thresholding removes all the Blue^-^ Red^-^ cells. All the Blue^-^ Red+ cells have Timer-Angle value 90, while all the Blue^+^ Red^-^ have Timer-Angle value 0. **(D)** Calculation of Timer Angle based on normalised Blue and Red fluorescence. **(E)** Timer locus model applied to a theoretical flow cytometric plot of Blue vs Red Timer fluorescence. **(F)** The 5 Timer loci are designated as follows: New = 0°; New-Persistent transition (NP-t) = (0°, 30°); Persistent [30°, 60°); Persistent-Arrested transition (PA-t) [60°, 90°); and Arrested = 90°

**Supplementary Table 1.** Differences between Tocky, Fate-mapper, and GFP/FP reporters.

|  | **GFP reporter (and other conventional FP reporters)** | **Fate-mapping reporter (e.g. Cre: Rosa-FP double transgenic)** |
| --- | --- | --- |
| **Time domain analysis** | No. Do not distinguish new expressors from pre-existing expressors | NA |
| **Frequency domain analysis** | No, due to long half-life of reporter proteins | No information |

| Tocky |
| --- |
| Reveal the time frame of early differentiation following key signalling |
| Identification of cells with different frequencies of transcriptional activity and finding unique functional states (e.g. antigen-specific T cells) |

